# A Novel Pilus System in Candidate Phyla Radiation Bacteria

**DOI:** 10.64898/2026.03.03.709456

**Authors:** Luca Troman, Junhyeong Kim, Jayson J. A. Rose, Matthew Johnson, Jillian F. Banfield, Steve Petrovski, Debnath Ghosal

## Abstract

The Candidate Phyla Radiation (CPR) represents a bacterial superphylum estimated to include between 15–50% of all bacterial species, yet CPR bacteria remain challenging to culture and have been primarily identified through metagenomic approaches. *Candidatus* Mycosynbacter amalyticus is a parasitic CPR bacterium that specifically targets *Gordonia amarae*, an actinobacterium with a hydrophobic, mycolic acid-rich cell envelope. Previous cryo-electron tomography indicated that *Ca*. M. amalyticus assembles thin extracellular filaments that are important for host interaction, yet their molecular identity remains unknown. Here, we applied single-particle cryo–electron microscopy to determine high-resolution structures of these filaments (2.8 and 3.6 Å), enabling the unambiguous identification of two previously uncharacterized pilins from the experimental density maps. These pilins, designated PamA and PamB, assemble into unique helical filaments distinct from all previously characterized filaments in both domain architecture and assembly mechanism. Despite low sequence identity, both PamA and PamB share conserved structural principles, including Ig-like folds and donor-strand exchange-mediated assembly. Phylogenetic analysis indicates that Pam pilins are exclusive to CPR bacteria, with homologues distributed predominantly across the classes Saccharimonadia and Microgenomatia. Analysis of the conserved *pam* operon identifies putative chaperones (PamC and PamD) and assembly factors structurally homologous to chaperone-usher pili components, suggesting an analogous but distinct assembly pathway. These findings expand the known diversity of bacterial pilus systems and demonstrate the power of structural approaches for characterizing uncharacterized proteins encoded within CPR genomes.

## Introduction

The advancement of high-throughput sequencing and genome-resolved metagenomics led to the recognition of Candidate Phylum Radiation (CPR) bacteria and prediction of their symbiotic lifestyles (1, 2). CPR represents a new frontier of microbial life; phylogenetic analyses of known CPR bacteria indicate that it constitutes the most genetically diverse superphylum currently known (3). CPR genomes harbor a plethora of unknown, uncharacterized, proteins, some of which may play critical roles in host association (4). Although members of the CPR are regularly detected in metagenomics, their obligate symbiotic lifestyle and specialized growth conditions make them very difficult to characterize. CPR have been identified from many specific environments including groundwater, soil, lake water, wastewater, and within human and insect microbiomes (5–8); however, relatively few examples of CPR have been cultivated in the laboratory due to the specificity of their environmental niches and lifestyles (7). Currently, all known CPR are obligate symbionts and a subset (Saccharibacteria) have been shown to associate with a specific actinobacterial host organism. Their parasitic lifestyle is predicted to compensate for their highly reduced metabolic capacity (9). Due to the absence of established genetic tools, molecular biology methods have rarely been applied to characterize the unique biology of CPR (9).

Previously, we characterized a Saccharibacterium, a member of the CPR group, from wastewater, *Ca.* M. amalyticus (henceforth referred to as M. amalyticus), and its interaction with host organism, *Gordonia amarae* (7, 10). Cryo-electron tomography (cryo-ET) of M. amalyticus and *G. amarae* cocultures uncovered an intimate surface interaction and the ultrastructural basis of M. amalyticus parasitism (10). The interaction involved M. amalyticus extracellular filaments, which attached to the host cell; and a tube-like structure that could facilitate interaction between the two microbes. Despite these advancements, we are yet to fully understand the mechanisms of host recognition and predation.

In our previous study (10), we found that the diameter of the thin filaments was consistent with those previously reported for type IV pili (11). Genes for Type IV pili have been identified with near universal abundance within CPR genomes (12), including type IV pilin gene annotation within the M. amalyticus genome (7), making this a plausible candidate. However, beyond canonical Type IV pili, bacteria produce diverse extracellular pili assembled from small pilin subunits with distinct architectures, including chaperone-usher pathway pili found within gram-negative bacteria and sortase-mediated assembly pili within gram-positive bacteria (13, 14). Although these systems differ in biological function, cellular context and assembly pathways, several pilins share common structural principles, including compact Ig-like folds and polymerization mediated by donor-strand exchange (15). Such pilin-like architectures recur across phylogenetically distant bacteria despite extensive sequence divergence, complicating their identification by sequence-based annotation alone (17–19). Given the evolutionary distance of CPR bacteria from well-characterized model organisms, and the lack of annotated CPR proteins, bioinformatic or proteomic approaches alone are unlikely to resolve the identity of the filaments that mediate their host interaction. In contrast, structural approaches such as cryo-electron microscopy (cryo-EM) can identify proteins directly from experimental density maps, bypassing the need for sequence homology or prior annotation (17–19).

Here, we employ single-particle cryo-EM to determine the structures of CPR pili at near-atomic resolution, enabling a “visual proteomics” strategy for protein identification directly from experimental density maps. Using this approach, we identify two previously uncharacterized pilins that are structurally distinct from any known pilus systems. We integrate these structural data with bioinformatic analyses, including structure prediction, sequence conservation, and gene neighborhood context, to propose models for pilus assembly and function. Together, our findings expand the known diversity of pilin (and bacterial filament) systems and provide new insights into the evolution and functional specialization of extracellular filaments in CPR bacteria.

## Results

### Two novel pilins identified within M. amalyticus pili

Isolated M. amalyticus were imaged using cryo-ET (Figure 1A and B). The resulting three-dimensional reconstructions and 3D segmentation showed multiple thin filaments originating from the cell body consistent with previous depictions of the M. amalyticus in co-culture with its prey, *G. amarae* (7, 10). Additionally, the sample contained several loose pili that appeared untethered from the M. amalyticus cell body (Figure S1). To investigate these filaments further, we used low-speed centrifugation to pellet intact cells whilst leaving detached pili in the supernatant. We then concentrated the soluble filaments within the supernatant and vitrified the sample on a cryo-EM grid for analysis by single-particle cryo-EM (Figure 1C). The filaments appeared uniform in the micrographs (Figure 1D). However, closer inspection of 2D class averages revealed two distinct pilus architectures (Figure 1E,I). We resolved each to sufficient resolution (2.8 and 3.6 Å) to predict the amino acid composition directly from the 3D electrostatic potential maps (Figure 1E-L, Figure S2). By searching against a theoretical proteome for M. amalyticus (generated from the genome NZ_CP045921.1), we used the automated model building software ModelAngelo (20) to generate initial molecular models for our experimental data (Figure 1G,K, Figure S3). This visual proteomics approach revealed the identity of each pilin class as distinct assemblies of two previously uncharacterized proteins (WP_260763287.1 and WP_260763280.1). The full genome-derived amino acid sequences for each pilus contained an N-terminal signal peptide, indicating SecY-dependent translocation through the bacterial inner membrane, consistent with their extracellular secretion. Importantly, mass spectrometry of cultured M. amalyticus confirmed that the two pilins were the 2^nd^ and 9^th^ most abundant proteins in whole-cell samples (Table S1). This not only validated our assignments but also demonstrated their abundance and indicated a central functional role for these unique pilins in M. amalyticus.

**Figure 1.**
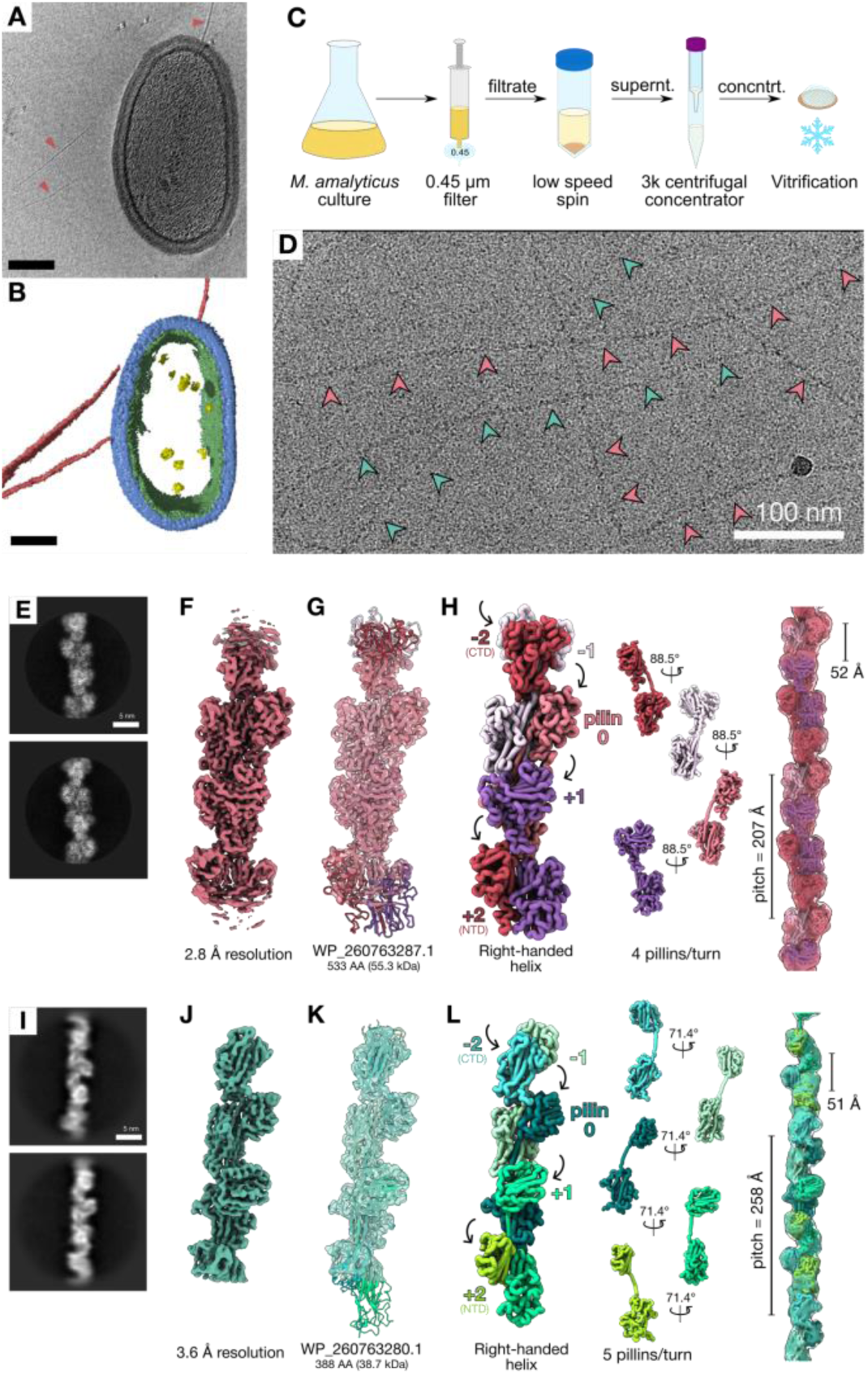
Identification of two novel pilins. (A-B) cryo-ET of isolated M. amalyticus: (A) tomogram slice through showing multiple thin filaments extending from the cell body (red arrowheads). (B) Segementation of the same tomogram. Pink – filaments; Green – inner membrane; blue – outer mycolic acid layer; yellow – ribosomes; Scale, 100 nm. (C) Schematic representation of the sample preparation workflow. M. amalyticus culture was filtered and centrifuged. The supernatent containing the detached pili was concentrated using a 3 kDa centrifugal concentrator and vitrified on a grid for cryo-EM analysis. (D) Representative cryo-EM micrograph showing morphologically uniform filaments. Red and green arrowheads indicate the two distinct pilus types identified by 2D classification. Scale bar, 100 nm. (E) Reference-free 2D class averages of the thicker pilus; (F) Cryo-EM reconstruction - estimated global resolution of 2.8 Å; (G) Atomic model of the pilin (WP_260763287.1) built into the cryo-EM density; and (H) Filament model – four pilin per helical turn (red, light pink, purple and rose) and right-handed helical twist of 88.5° and pitch of 207 Å. (I) Reference-free 2D class averages of the thinner pilus; (J) Cryo-EM reconstruction with estimated global resolution of 3.6 Å; (K) Atomic model of the pilin (WP_260763280.1) built into the cryo-EM density; (L) Filament model – five pilins per helical turn (cyan, pastel green, dark green, light turquoise, light green) and right-handed helical twist of 71.4° and a pitch of 258 Å.

Both pili shared several structural features; both comprised of two domains assembled into elementary right-handed helices with helical rises of 51–52 Å (Figure 1H and L). However, the two classes differed both in width and helical parameters. The thicker pilus comprises larger pilins (55.3 kDa) with a greater twist angle (88.5°) and lower helical pitch (207 Å) (Figure 1H). In contrast, the thinner pilus is assembled from smaller pilins (38.7 kDa) with a reduced twist (71.4°) and extended pitch (258 Å), resulting in a visibly looser helical arrangement (Figure 1L). Despite these differences, distinguishing the two pilus types directly from raw cryo-EM micrographs proved challenging due to inherently low signal-to-noise ratio (Figure 1D), but the ability to computationally classify the two filaments apart indicates compositionally homogeneity for long stretches within each filament.

To investigate whether the two pilins formed separate filaments or were incorporated sequentially within the same filament, we overlaid particle coordinates from the final refinements of each pilus class back onto the original micrographs (Figure S4). We observed multiple clear instances where contiguous stretches of particles assigned to PamA switched to stretches assigned to PamB within a single filament consistent with the sequential incorporation of different pilin subunits into a single filament (Figure S4).

### Both pilins share a domain architecture and assembly

Analysis of the domain architecture showed that both pilins comprise N- and C-terminal domains linked by a single β-strand (Figure 2A-C and F-H). In both cases, the N-terminal domains (NTDs) contain nine β-strands arranged in a β-sandwich. However, the larger of the two pilins includes ∼100 additional amino acids that form two additional β-strands and two short α-helices compared with the smaller pilin (Figure 2A-C and F-H).

**Figure 2.**
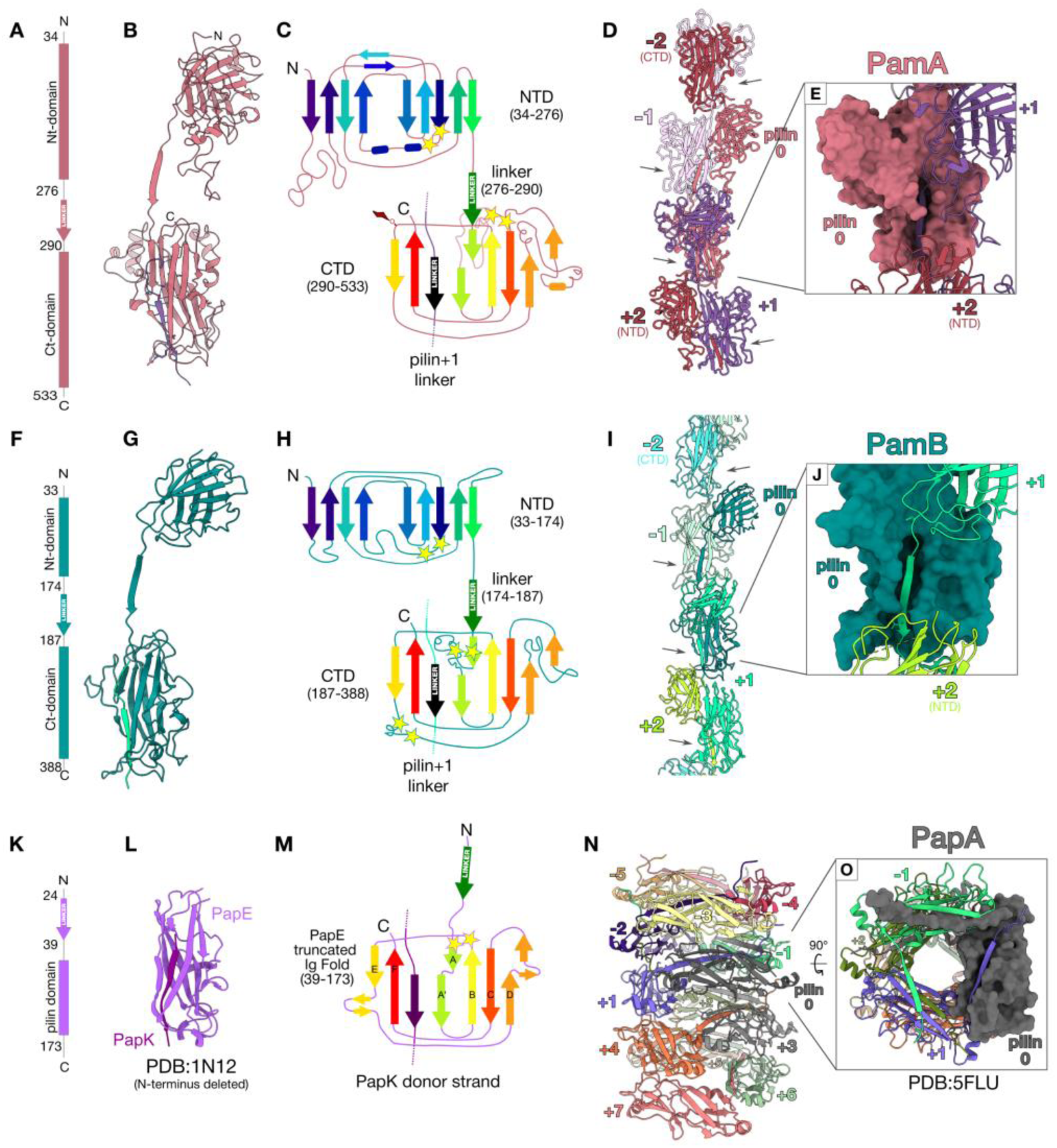
Domain architecture and assembly of Pam Pilins. (A-E) PamA pilin. (A) Cartoon domain architecture, (B) cryo-EM derived molecular model, (C) secondary structures cartoon depictions, (D) assembled pilus model with (E) close-up of donor strand (cartoon) within the CTD from the adjacent pilin (surface). (F-J) PamB pilin. (F) Cartoon domain architecture, (G) cryo-EM derived molecular model, (H) secondary structures cartoon depictions, (I) assembled pilus model with (J) close-up of donor strand (cartoon) within the CTD from the adjacent pilin (surface). (K-M) Typical truncated Ig-like domain. (K) PapE cartoon domain architecture, (L) Crystal structure of PapE-PapK (PDB:1N12) (23) in cartoon depiction, (M) PapE secondary structures depiction with PapK donor strand. (M-N) cryo-EM structure of PapA assembly in P-pilus (PDB:5FLU).

The C-terminal domains (CTDs) of both pilins are more similar in size and overall fold, forming an open β-barrel that accommodates the linker strand of the adjacent pilin within the filament (Figure 2D-E and I-J). Interestingly, the CTD fold and its interaction with the linker closely resembled the single truncated Ig-like domain described within chaperone-usher pilin systems (15, 21) (Figure 2K-M) (22). Sequence comparisons across other closely related pilins within other CPR bacteria show that in M. amalyticus pilins, the donated linker and CTD binding site are the most highly conserved regions (Figure S5), consistent with a conserved strand-exchange-based assembly mechanism. Given the similarities to gram-negative the Pap/Fim pilus system, we term the M. amalyticus pilins “Pam” and our two pilins PamA and PamB.

Despite these parallels, Pam pili differ from Pap/Fim pili systems in terms of domain architecture, pilus assembly, and length. For example, across both Pap and Fim pilus systems, the major pilins (PapA and FimA) coil in on themselves in a helical arrangement with three pilins per turn and a helical pitch of 25.2 Å (Figure 2N-O). In these extracellular matrix pili, the arrangement of the major pilins allows coiling and uncoiling of the filament, giving remarkable dynamic and expandable properties that allow the maintenance of adhesion within the eukaryotic host (23). Within the coiled PapA pilus (PDB:5FLU) (23), the interface area between two adjacent pili with the donated β-strand is estimated to be 1500 Å^2^. However, this is further strengthened by additional interfaces between proximal neighboring pili within the coil, thus a total interaction interface of 2458 Å^2^ per pilin is predicted within the coiled filament.

In Pam pili, both pilins form elementary right-handed helices, where coiling is sterically hindered by the additional domain. In place of coiling, the NTD contributes to the interface areas between M. amalyticus pili. For the thicker filament, named PamA, the interface between adjacent pilins extends to 2584.7 Å^2^ with a predicted solvation free energy gain (ΔG_solvation_) of -26 kcal/mol and intermolecular formation of 37 hydrogen bonds and 4 salt bridges. For the thinner filament, PamB, the interface area between adjacent pilins was 1907.3 Å^2^ with a predicted solvation free energy gain (ΔG_solvation_) of -18.5 kcal/mol and intermolecular formation of 22 hydrogen bonds and four salt bridges. Additionally, there is a small interface between one pilin and the next furthest pilin (n+2/n-2) of 325.6 Å^2^ in PamA and 319.5 Å^2^ in PamB. Therefore, the total interfaces for an incoming pilin with the existing filament extend to 2910.2 Å^2^ in PamA and 2226.8 Å^2^ in PamB. These are within the same range as the coiled PapA pilins (2458 Å^2^).

Additionally, we identified two and three sets of paired cysteines within PamA and PamB pilins, respectively (yellow stars, Figures 2C and H), with the potential to form disulfide bonds that may enhance filament stability and rigidity. Finally, at the resolution obtained for the PamA filament, we confidently identified additional density bound to Serine356 of PamA, consistent with post-translational modification (PTM) (Figure S6). Taken together, the extensive interaction interface, intermolecular bonding, PTM and disulfide stabilization reflect adaptations to the extracellular environment in which these bacterial filaments function.

### Phylogenetic survey points to CPR-specific evolution of PamA and PamB

To contextualize these findings, we surveyed the phylogenetic distribution of PamA and PamB across bacterial genomes using the GTDB-CPR database (Figure 3). Both proteins were detected exclusively within CPR bacteria, with no homologues identified in non-CPR lineages, indicating that Pam pili are specific and unique to this superphylum. The highest prevalence of Pam pilins were observed in the classes Saccharimonadia and Microgenomatia (Figure 3). Using a pure MMseqs2 search, we recovered 88 genomes encoding PamA homologs and 344 genomes containing PamB homologs predominantly restricted to Saccharimonadia. We then used a more sensitive language model–guided search to identify homologs that might be missed by sequence identity alone (as has been previously observed (25)), this expanded this set by 304 additional PamA-positive genomes. We compared predicted protein structures of a Microgenomatia homolog identified by the language-model searches from Candidatus Roizmanbacteria bacterium and this appeared to have conserved structural homology, despite the low sequence identity (20.95% similarity to M. amalyticus PamA) (Figure S7). This is consistent with the low sequence identity to other bacterial pilins which share the conserved Ig-like folds and donor strand assembly mechanisms and implies that sequence-based homology searching underestimates conservation within CPR, and Pam pili could be even more widespread than showed within this data.

**Figure 3.**
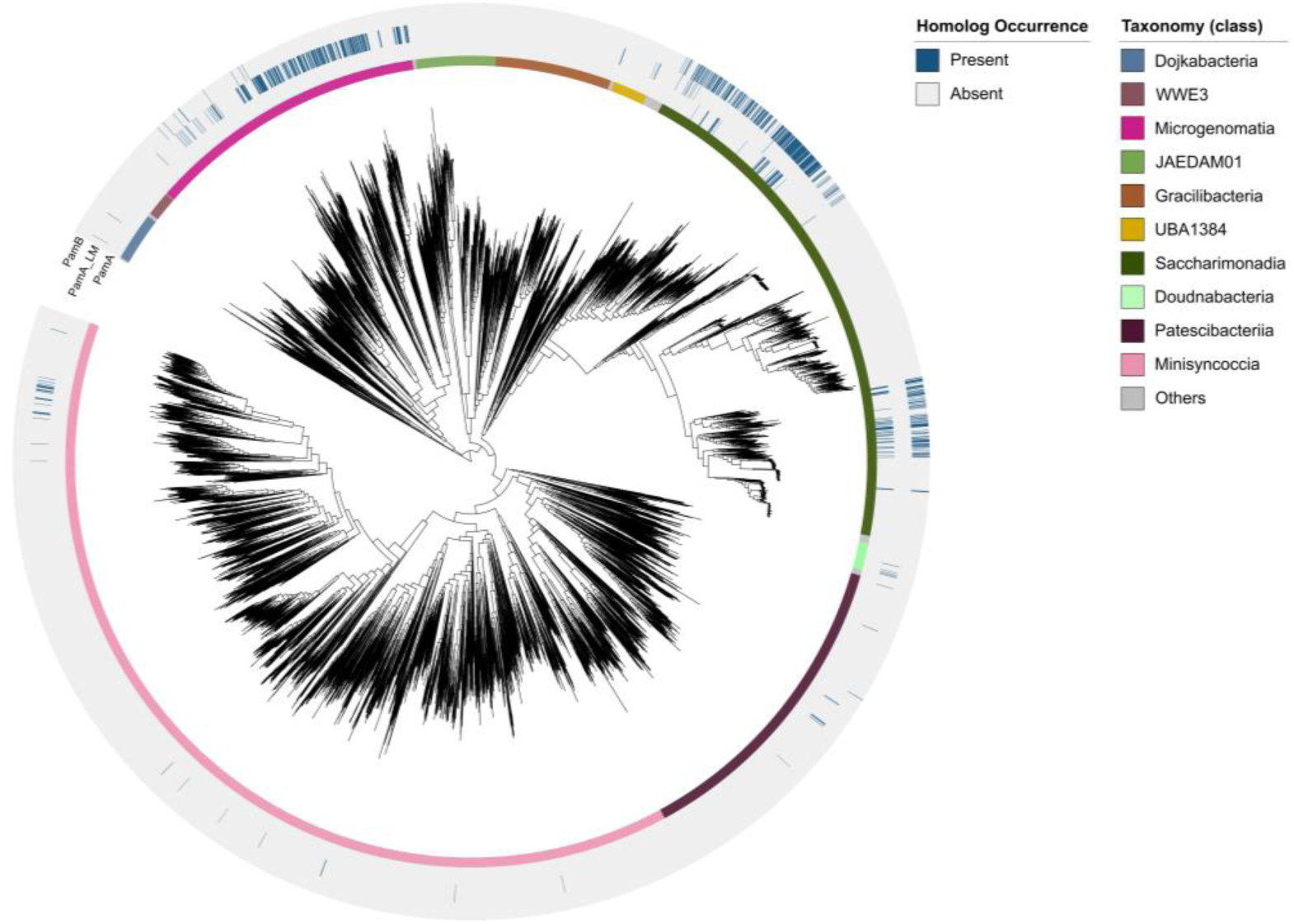
PamA and PamB distribution across CPR genomes. Circular phylogenetic tree of CPR bacteria constructed from the GTDB-CPR database, overlaid with occurrence patterns for PamA and PamB homologs. PamA and PamB homologs were identified using MMseqs2 with an e-value threshold of 10⁻⁵. Language model–based searching recovered additional distant PamA homologs, designated PamA_LM, expanding the known distribution of PamA across CPR lineages. The outer ring displays homolog occurrence (present/absent) for each genome, whilst the inner ring shows taxonomic classification at the class level, revealing that Pam pilins are distributed across multiple CPR lineages with highest prevalence in Saccharimonadia and Microgenomatia.

### Pilin assembly mechanism

Within Pap/Fim filaments in gram-negative bacteria, pilin assembly components are all encoded within the same operon. Similarly, we found both Pam pilins are encoded within the same operon in M. amalyticus alongside several other uncharacterized proteins, we named this the *pam* operon (Figure 4A). To investigate their potential roles in pilus assembly, we used AlphaFold (25) structural predictions in combination with FoldSeek (26), to search for structural homologues that might provide mechanistic insights into assembly and function (Figure 4B).

**Figure 4.**
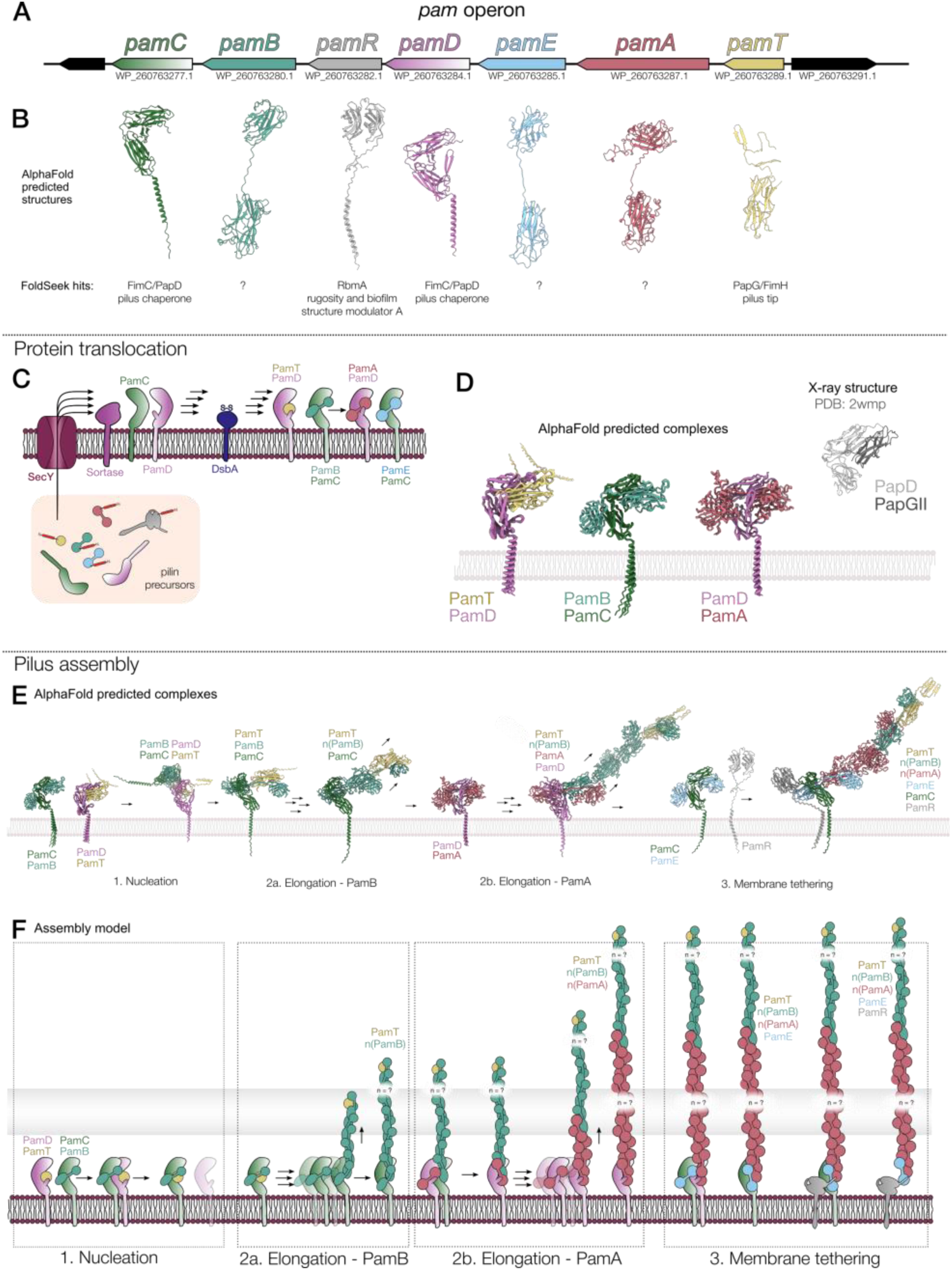
Pam pilus assembly mechanism and predicted protein complexes. (A) *pam* operon in M. amalyticus. (B) AlphaFold-predicted structures of *pam* operon components (with N-terminal signal peptides omitted) with structural homologues identified by FoldSeek for each protein. (C) Predicted protein translocation pathway. Pilin precursors (shown in inset) are translocated across the inner membrane via SecY in a signal peptide-dependent manner. Membrane-anchored chaperones (PamC and PamD) facilitate donor-strand exchange and stabilize pilin intermediates during assembly. (D) AlphaFold predicted Pam chaperone-pilin complexes and the X-ray crystal structure of the PapD-PapGII complex (PDB: 2wmp)(28) for comparison. (E) Pam complexes predicted by AlphaFold indicate an ordered Pam pilus assembly pathway. (F) Schematic of our model: pili are nucleated by PamT (tip pilin) at the distal end, followed by elongation by the sequential addition of PamB, PamA, and PamE pilins. Membrane-anchored chaperones (PamC and PamD) facilitate each donor-strand exchange step. The RbmA-like protein PamR terminates elongation and anchors the completed filament to the inner membrane.

Signal peptide analysis (SignalP - 6.0 (25)) identified N-terminal signal sequences in all components, consistent with SecY-mediated secretion into the bacterial envelope. Structural predictions for WP_260763277.1 and WP_260763284.1 revealed homology to the FimC and PapD periplasmic chaperones, which govern Pap/Fim filament assembly in gram-negative bacteria (Figure 4B). Consequently, we named these PamC and PamD, respectively. Unlike canonical Pap/Fim chaperones, PamC and PamD both contain a C-terminal transmembrane helix, indicating that they remain anchored in the inner membrane after SecY translocation, rather than diffusing freely in the periplasm. PamC and PamD differ at the hinge region between the two β-barrel domains; here PamD an additional β-hairpin relative to PamC (Figure 4B).

Several other operon-encoded proteins had predicted folds compatible with the donor strand exchange mechanism proposed for pilus assembly (Figure 4B, Figure S7). WP_26076385.1, we assigned as PamE, exhibited a domain architecture highly similar to the cryo-EM resolved pilins PamA and PamB, characterized by the N-terminal nine-strand β-sandwich, a linker strand, and a C-terminal truncated Ig-like domain. Another protein, the single domain WP_260763289.1, contained only a truncated Ig-like domain but lacked a linker strand. This architecture resembles the tip pilins in Pap/Fim systems and thus may represent the initial donor-strand acceptor at the pilus tip – we therefore named this PamT.

Additionally, WP_260763282.1 had a predicted fold resembling the extracellular matrix scaffolding protein RbmA (Rugosity and Biofilm structure Modulator A) from *Vibrio cholerae*, which contains two tandem fibronectin type III (FnIII) domains. In *V. cholerae*, RbmA functions as a linker between the cell surface and extracellular polysaccharides (EPS), stabilizing biofilm architecture. Our predicted RbmA-like protein – now renamed PamR - also contains a C-terminal transmembrane helix and an additional N-terminal β-strand, features that may tether the pilus directly to the membrane. Taken together, these similarities and differences suggest an analogous role in Pam pilus attachment.

To further probe the assembly mechanism, we used AlphaFold (25) to predict complexes between the putative chaperones and pilins (Figure 4D, S8). The predicted chaperone-pilin interactions mirrored known Pap chaperone-pilus complexes from *E.coli* (PDB:2wmp) (28). Specifically, β-strands from the chaperone were predicted to insert into the open β-barrel of the pilin CTD, stabilizing its truncated Ig-fold prior to donor strand exchange with the incoming pilin. Based on these models, we propose that Pam chaperones facilitate donor strand exchange to drive filament elongation, while their transmembrane helices anchor the complex to the inner membrane in the absence of an outer membrane.

Larger predicted assemblies of chaperone-pilin complexes suggested an ordered filament assembly pathway (Figure 4E, S9). PamT, the single domain pilin, was predicted to occupy the tip position, with the linker from the next pilin occupying the Ig-like domain. Complex predictions indicated a likely assembly sequence: PamT at the tip, followed by a stretch of either, or a combination of PamB, PamA, and finally PamE. Incorporation of the RbmA-like PamR into the models revealed its N-terminal insertion into PamE, suggesting a role in terminating pilus elongation and tethering the structure to the membrane.

Taken together, our analysis support three conclusions (i) there are no specialized transenvelope assembly machinery - instead, donor strand exchange mediated by membrane-anchored chaperones is sufficient for filament growth, (ii) our mass spectrometry and cryo-EM data indicate long stretches of PamA and PamB which represent compositional domains within a single pilus (iii) the RbmA-like PamR likely terminates pilus assembly and anchors the filament to the inner membrane.

### Neighborhood analysis reveals conserved gene context surrounding PamA

To understand the genetic context and conservation of the *pam operon*, we compared the gene neighborhood of the M. amalyticus *pam* operon with other CPR bacteria that encode the PamA homologs shown in Figure 3. Analysis of 30 kb genomic windows centered on *pamA* revealed a strongly conserved gene context, including the putative *pam* operon (Figure 5). A sequence similarity network (SSN) constructed from these genomic windows confirmed the conservation of core *pam* operon components across CPR genomes (Figure 5A). Within this network, genes encoding ATP synthase subunit proteins and a tail fibre–like protein occupied central positions, indicating their consistent co-localization with the *pam* operon across diverse CPR lineages.

**Figure 5.**
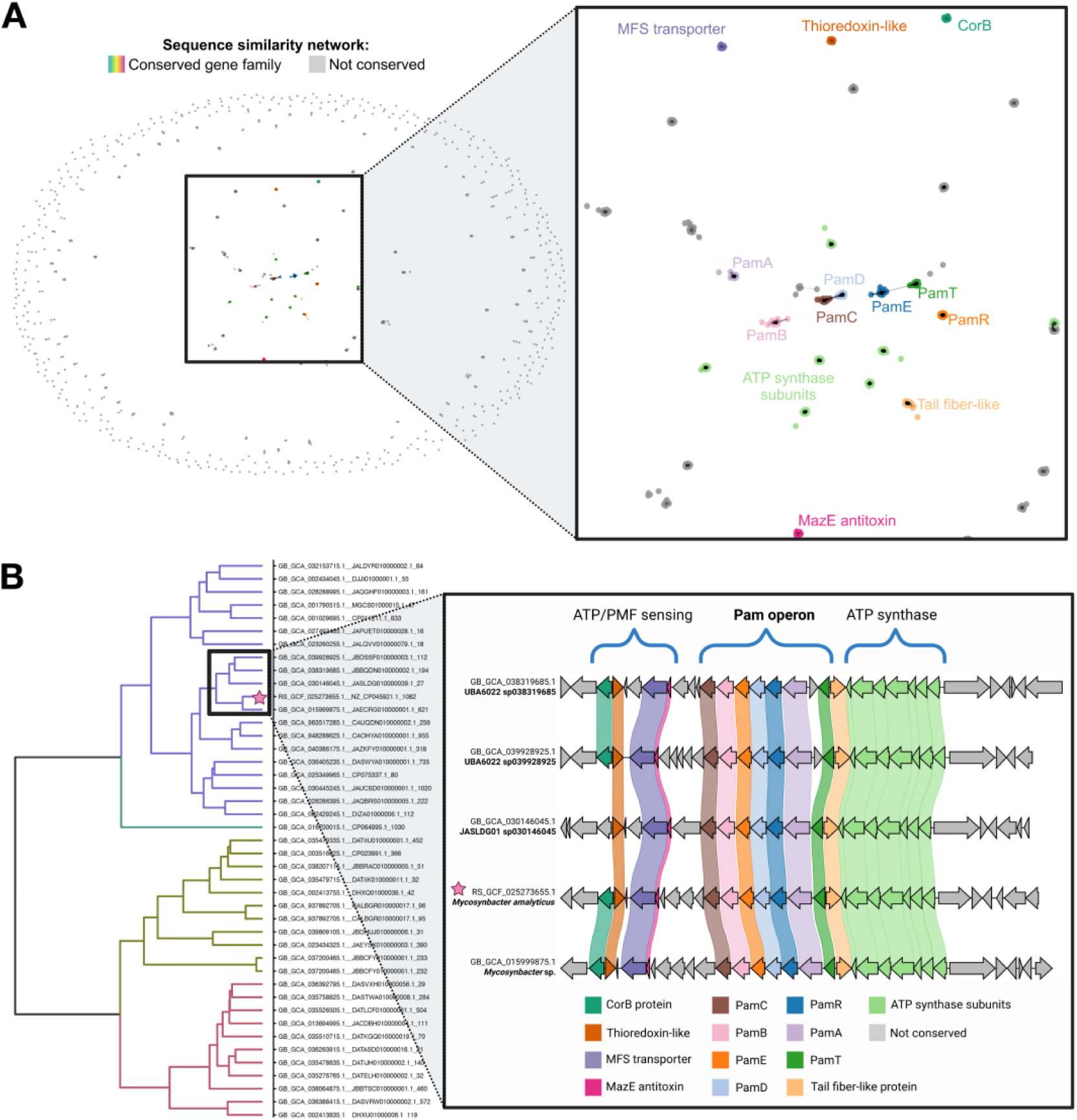
PamA neighbourhood analysis. (A) Sequence similarity network (SSN) analysis of 30 kb genomic windows centered on PamA across CPR genomes. Each node represents a gene family, with node size indicating prevalence and edges indicating sequence similarity. Genes not conserved across the dataset are shown in grey. The inset (right) provides a zoomed view of the central Pam operon region. (B) Dendogram (c-blaster based) showing PamA neighborhood similarity across CPR lineages. Left: Phylogenetic tree of CPR bacteria containing PamA homologs, with major clades indicated by colored branches. The M. amalyticus JR1 strain is marked with a star and highlighted with a black box. Right: Gene cluster diagrams showing the genomic organization of the *pam* operon and flanking regions across representative genomes from different CPR lineages. Three conserved regions are indicated: ATP/PMF sensing (left), the core *pam* operon (centre), and ATP synthase components (right). Gene orientation and color indicate orthology relationships.

Analysis of neighborhoods most closely related to the M. amalyticus strain JR1, revealed conserved genes predicted to function in ATP-sensitive and proton motive force (PMF)–driven transport (Figure 5B). Specifically, the M. amalyticus gene annotated as *corB* (WP_260763263.1) encodes a protein predicted to contain a cytoplasmic ATP-binding domain with structural architecture similar to archaeal CorB (Figure S10A). Within Archaea, CorB dimerises to form an ATP-sensitive Mg^2+^ transporter, an essential regulator of Mg^2+^ homeostasis (29). Additionally, an adjacent gene (WP_260763267.1) encodes a membrane protein homologous to major facilitator superfamily (MFS) transporters, which couple the proton motive force to substrate transport across the inner membrane (Figure S10B). This MFS transporter gene is located immediately upstream of a gene encoding a MazE-like antitoxin that neutralizes the nucleic acid-binding toxin MazF, however, only the MazE-like antitoxin is encoded here. The conserved association of these components with the Pam operon suggests functional integration with pilus machinery, though the precise mechanisms remain to be determined.

## Discussion

Here, we have used a “visual-proteomics” approach to identify unknown bacterial pilins directly from experimental cryo-EM data. The application of several recent advances in cryo-EM acquisition, data processing, and deep-learning-based molecular model building software has made this possible. Using these advanced methods, we describe a novel filament system in CPR bacteria which is distinct from any previously characterized bacterial filament system. The unique nature of these pilins obscured their identification through conventional sequence-based approaches. This was in part due to the lack of any meaningful genomic annotations that traditionally could have enabled their identification. This highlights the strength of this approach for studying the vast repertoire of uncharacterized proteins encoded within CPR genomes. Our findings expand our understanding of molecular complexity and evolutionary innovation within this superphylum, revealing adaptations underlying host associated lifestyles that remain hidden from conventional genomic annotation.

### Streamlined linear pilin assembly promotes the reach of Pam pili

In the case of our novel Pam filaments, we found several similarities to the canonical Pap/Fim pili found within gram-negative bacteria. Like Pap/Fim, the Pam pili are assembled by donor-strand exchange with extensive intermolecular interfaces; pilins contain several intramolecular disulphide bonds that stabilize their assembly. However, the differences are substantial. Pap/Fim filaments comprise a single major pilin that coils in on itself to form a dynamic and expandable coiled helix, with multiple contacts between proximal neighbors within the coiled arrangement. In contrast, Pam pili are assembled from two major pilin classes (PamA and PamB) that remain in an extended, linear assembly. In Pam pili, the NTD compensates for the absence of coiling, providing additional intermolecular contacts that stabilise the linear filament. Thus, despite this architectural difference, both systems achieve comparable intermolecular interface areas per pilin (∼2458 Å² for Pap versus 2910.2 Å² for PamA and 2226.8 Å² for PamB).

We propose that these architectural differences reflect distinct functional strategies. Pap/Fim pili mediate dynamic cell-cell adhesion through coiling and uncoiling, allowing bacteria to maintain stable yet flexible tethers to host cells. In contrast, Pam pili appear to be optimized for environmental reach: the linear geometry extends the filament length per pilin relative to coiled counterparts, reducing the metabolic cost of assembly whilst maximizing reach. This is consistent with our microscopy data (Figures 1A and S1) showing that M. amalyticus produce multiple pili extending far from the cell body. Combined with the observation that these pili appear easily untethered, we suggest that Pam filaments represent an adaptation for low-cost production of an extended filament net, a strategy fundamentally different from the dynamic tethering of gram-negative pili.

The two distinct pilin classes likely serve complementary capture functions. Our predicted modelling reproducibly suggest that PamB assembles at the distal end, followed by PamA at the proximal end to the cell body (Figure 4E). We hypothesise that extended PamB-type filaments extend furthest into the environment to detect and contact the mycolic acid-rich cell envelope of *G. amarae*, whilst the rigid PamA-type filaments form structures that provide strength at their base to maintain an interaction with the host.

### Establishing host contact

Our previous cryo-ET work revealed that *G. amarae* predation depends on recognition of the mycolic acid-rich outer layer by thin filaments extending from the M. amalyticus cell body (10). We initially hypothesised these filaments might represent retractable Type IV pili. However, the Pam pilin structures we have now resolved differ substantially from canonical Type IV pili in both architecture and assembly, indicating a distinct interaction mechanism consistent with a parasitic lifestyle.

The expansive reach of the Pam filaments indicate that they could establish initial contact with the host cell. However, several critical questions remain unresolved. First, we do not know how pilins interact with the mycolic acid layer or whether post-translational modifications enhance recognition. Second, we could not identify retraction machinery within cryo-ET data or the gene neighbourhood, leaving unresolved whether Pam filaments serve a passive role in establishing association or if they are actively retracted following adhesion to facilitate host cell capture. Lastly, we do not know if or how these Pam filaments might cooperate with other filamentous/tube-like structures identified within M. amalyticus (29). These mechanistic details will require further structural and functional investigation.

Functional specialisation of distinct filament systems is not unique to M. amalyticus. Recent structural studies of other Saccharibacteria (TM7 CPR bacteria) (30) identified two functionally distinct Type IV pili: a thick, retractable pilus mediating twitching motility; and a thin, non-retractable pilus involved in host binding and episymbiotic growth. Similarly, contact-dependent predation systems employ multiple filament types for prey capture and killing (31). These observations across diverse parasitic systems suggest that functional specialisation of filament systems represents a conserved host interaction strategy across diverse bacterial lineages.

### Gene neighbourhood indicates coupling of pilus assembly to M. amalyticus metabolism

The prominent co-localisation of ATP synthase components within the *pam* operon neighbourhood across diverse CPR bacteria indicates a conserved association of pilus assembly machinery with ATP-production. Additionally, genes encoding ATP or PMF-driven substrate transporters (including a CorB-like protein and a predicted MFS-family protein) are conserved within the *pam* operon neighbourhood in close relatives of M. amalyticus. These observations suggest possible functional integration of pilus machinery with cellular metabolism, warranting further investigation.

This study demonstrates the power of integrating state-of-the art cryo-EM with *in silico* structural biology to assign molecular identity and function to previously uncharacterised proteins – without reliance on sequence homology or prior annotation. Our current study holds broad relevance for understanding of CPR episymbiosis more widely, given the large diversity of CPR bacteria that encode the *pam* operon and the conserved coupling of pilus assembly with metabolic components. The structural parallels between Pam and Pap/Fim pilins, despite low sequence identity, raises intriguing questions about the evolutionary origins of these systems. This work provides both a methodological blueprint for illuminating the biology of CPR and other poorly characterised lineages that have resisted conventional investigation, and a structural foundation upon which future genetic and functional studies can build.

## Acknowledgements

This project was supported by an NHMRC grant (APP1196924), an HFSP grant (RGEC33/2023) and the Cumming Global Centre for Pandemic Therapeutics, Peter Doherty Institute for Infection and Immunity grant (CGCPT00060) awarded to DG. J.J.A.R was the awardee of a La Trobe University postgraduate award. This work was funded by university funds provided by the school of agriculture biomedicine and environment to S.P. We also acknowledge the Ian Holmes Imaging Centre at the University of Melbourne for access to microscopy facilities. Cryo-EM data processing was supported by The University of Melbourne’s Research Computing Services and the Petascale Campus Initiative. We would also like to thank Rohan Lowe and the La Trobe University Proteomics and Metabolomics Platform for the provision of instrumentation, training and technical support in acquisition of proteomics data. Support was also provided by the National Institute of Allergy and Infectious Diseases of the National Institutes of Health under award number 5R01AI092531 and the Emerson Collective.

## Data availability

Cryo-EM reconstructions of the PamA and PamB filaments are deposited under EMD-80296 and EMD-80297 respectively; corresponding atomic models are available in the Protein Data Bank under accession codes PDB 25PZ and 25QA. Custom scripting has been made publically available through deposition to github (github.com/Luca-Atlas/Micrograph_sorting). The mass spectrometry proteomics data have been deposited to the ProteomeXchange Consortium via the PRIDE partner repository with the dataset identifier PXD078107.

## Author Contributions

Conceptualization, D.G., S.P.; study design, D.G., S.P., L.T., J.J.A.R.; methodology, L.T., J.J.A.R.,

M.J., J.K., J.F.B., S.P., D.G.; formal analysis, L.T., M.J., J.K.; investigation, L.T., J.J.A.R., M.J., J.K.; data curation, L.T.; writing – original draft, L.T., M.J., J.K.; writing – review & editing, all authors; funding acquisition, S.P., J.F.B., D.G.

## Materials and Methods

### Bacterial strains and growth

Parasitic M. amalyticus strain JR1 cells were cocultured with *G. amarae* CON44^T^ host cells as previously described (7, 10). Briefly, 300µL of M. amalyticus was flooded onto and cocultured on *G. amarae* CON44^T^ lawns and incubated on solidified PYCa (5 g.L^-1^ peptone, 3 g.L^-1^ yeast extract, 1 g.L^-1^ calcium chloride and 1 g.L^-1^ glucose) agar plates (12 g.L^-1^ agar) at 28 °C for 48 h.

M. amalyticus cells harvested for cryo-EM were resuspended in 3mL of PYCa broth using a sterile glass rod, with resulting suspension filtered using a 0.45 µm filter (Sartorius).

### Proteomic extraction and mass spectrometry (MS) analysis of M. amalyticus cells

M. amalyticus cells were harvested from a CON44^T^ lawn plates with lysate resuspended with milliQ H_2_O replacing PYCa broth to minimise contaminating proteins present in the PYCa media and were filtered using a 0.22 µm filter (Sartorius). 4 mL of the harvested M. amalyticus cells were centrifuged at 21,380 x *g* for 1 hour, most of the supernatant was discarded, and the pellet was resuspended in 100 µL of retained supernatant. The resulting resuspension was then sonicated for two 30s bursts with a sonifier 250 (Branson). Protein concentration was then quantified using a Qubit 3 fluorometer (Invitrogen). Following extraction, protein was prepared for MS analysis using the single pot solid-phase-enhanced sample preparation (SP3) method (32). Briefly, 20 µg of protein sample were reduced and alkylated with 25 mM tris 2-carboxyethyl phosphine (TCEP) and 120 mM iodoacetamide (IAA), respectively. The alkylated sample then underwent bead clean up and digestion using Stock SpeedBeads carboxylate (Cytiva), and Trypsin in 20 mM ammonium bicarbonate, pH 8, respectively. Samples were then analysed with liquid chromatography-mass spectrometry (LC–MS) Orbitrap Eclipse (ThermoFisher). Peptides detected through LC-MS analysis were then compared to the hypothetical proteome of M. amalyticus (NZ_CP045921.1) and *G. amara*e (NZ_CP045810.1). All remnant host peptides that matched *G. amarae* were then filtered from the resulting data set. The remaining peptides of M. amalyticus origin were then sorted by normalized abundance, reflecting ∼75% coverage of the predicted M. amalyticus proteome. Genes encoded in the *pam* operon were then identified with their corresponding abundance.

### Cryo-electron tomography sample preparation

Planktonic cell suspensions of M. amalyticus were concentrated to an OD_600_ of 2.0 by centrifugation and mixed with 10 nm colloidal gold beads (precoated with BSA) (Sigma-Aldrich, Australia) in a ratio of 4:1. The sample was applied to a glow discharged copper Quantifoil holey carbon grids (Quantifoil Micro Tools GmbH, Jena, Germany). Blotting and vitrification were done using a Vitrobot IV (FEI Thermo Fisher Scientific) with chamber humidity set to 100%. Samples were blotted for 6-8 seconds with a blot force of 8 before rapid plunge vitrification in liquid ethane.

### Cryo-ET data collection

Grids were imaged using Titan Krios G4 cryo-EM operating at 300 kV acceleration voltage, and a K3 direct detector equipped with a Gatan bio quantum energy filter (slit width 20 eV), in CDS super-resolution movie mode. The tilt--series was acquired with a dose-symmetric tilt scheme using Tomography 5 software version 5.14 (Thermofisher Scientific) using a tilt range of ±54° with 3° increments. Data were collected with a total electron dose of 140 e^-^/Å^2^, a defocus of -6 μm, and a bin 2 pixel size of 3.39 Å/pixel.

### Cryo-ET data processing

Data processing was done using the following software packages within Scipion (33). Motion correction was done using MotionCor3 (34). Tilt-series alignment was with IMOD (version 5.1) using x4 binned tilt-series (13.56 Å/pixel), and tomogram reconstruction was done using Tomo3d. The tomogram was initially segmented using membrainV2 (35) to segment membranes, filaments and ribosomes were segmented using Dragonfly software (https://www.theobjects.com/dragonfly/index.html). Briefly, a neural network 5-class U-Net (with 2.5D input of 7 slices) was trained on tomogram slices to recognise M. amalyticus pili, and ribosomes. All segmented features were fine-tuned to achieve high-quality 3D segmentation. 2D images were created using Dragonfly.

### Filament isolation

1mL of the M. amalyticus culture (described above in bacterial strains and growth) was centrifuged at 21,380 x *g* for 15 minutes, retaining the filament containing supernatant and discarding the pelleted cells. Using a 3K Ultra-0.5 centrifugal filter device (Amicon), the resulting supernatant was concentrated (6X), as per manufacturer’s instructions. Briefly, 500 µL of supernatant was centrifuged at 14,000 x *g* for 15 minutes, followed by inversion of the filter device and a 1,000 x *g* centrifugation of the concentrated solute for 2 minutes.

### Single particle cryo-EM sample preparation

Shortly before use, Quantifoil R1.2/1.3 holey carbon grids (Quantifoil Micro Tools GmbH, Jena, Germany) were glow discharged. Grids were then transferred to the humidity-controlled chamber of a Leica EM GP2 (95% relative humidity, 4 °C). To increase sample concentration on the grid, two successive applications of 4 µL sample were performed, each followed by back blotting to remove excess liquid (sensor-enabled blotting, blot distance 1 mm, blot time 5s). Grids were immediately vitrified by plunge freezing into liquid ethane.

### Cryo-EM single particle data collection

High-resolution data for single-particle cryo-EM analysis were collected on FEI Titan Krios G4 300 keV FEG transmission electron microscope (Thermo Fisher Scientific). Micrographs were recorded using a K3 Summit direct electron detector (Gatan) equipped with a post-column energy filter set to a slit width of 10 eV. Data were acquired using automated EPU software at a nominal magnification of 105,000 × with pixel size of 0.834 Å and a defocus range of -0.5 to -2.4 µm.

### Cryo-EM single particle data processing

With the exception of particle picking, all image processing steps were carried out using RELION-5 (36, 37). Beam-induced motion within each movie was corrected using MotionCor2 (34), and the parameters of the contrast transfer function (CTF) for each motion-corrected, dose-weighted movie were determined using CTFFIND4 (38), both within the RELION interface. The resulting micrographs were imported into crYOLO (v1.9.9) (39, 40) for automated filament picking. Manually picked filaments from 45 micrographs were used to train an initial model to pick across a range of different defocus values. The model was used to pick 118,784 filaments from the 7,026 micrographs with intervals of 62 pixels between boxes (∼52 Å), minimum length of 4 particles, and an optimized threshold of 0.3.

The resulting coordinates were re-imported into RELION-5 with a box size of 320 × 320 pixels (267 × 267 Å) and 4× binning to 80 × 80 pixels. To roughly align particles to the helical axis 4× binned particles were subjected to supervised 3D classification for four iterations using a reference reconstructed from a small screening data set low pass filtered to 20 Å. 2D classification of the roughly aligned particles (with no image alignments) allowed selection of 53 classes of the major pilin containing 389,566 particles for further processing. Additionally, a 2D classification with image alignments allowed separation of a second class of helical filaments which were subsequently re-extracted with refined offsets reset to zero. For both pilus types, particles were further refined through 2D classification, iterations of CTF refinement and Bayesian Polishing. RELION post-processing of unsharpened half-maps from each refinement were used to generate Fourier Shell Correlation (FSC) analysis to describe the stated resolutions of 2.8 and 3.4 Å for each pilus. Finally, deepEMhancer (41) of the unsharpened half maps were used for local sharpening to improve the interpretability for model building.

### Protein identification, model generation and model refinement

The final post-processed reconstructions were analyzed using ModelAngelo (20) to search against a hypothetical proteome generated from open reading frames in the genome for the organism (NCBI reference sequence # NZ_CP045921.1). Custom scripting was used to process the output and identify the fragments. ModelAngelo was then rerun using a FASTA file containing the amino acid sequences for the putative pilins generating initial models for each chain that served as starting models for further refinement. Missing residues were built in Coot (42), and the model was subsequently refined using multiple iterations of ISOLDE (43) in ChimeraX (44) using interactive, force field-based molecular dynamics fitting against both the deepEMhancer (41) post-processed density and the unsharpened maps from RELION 3D auto-refinement, and Phenix real space refinement (45) and Coot (42) using only unsharpened maps.

### Model and conservation analysis

Protein–protein interfaces in the refined atomic models were analysed using PISA (Proteins, Interfaces, Structures and Assemblies) with default settings (46). Reported metrics include buried surface area (Å²), estimated solvation free-energy gain on complex formation (ΔG, kcal·mol⁻¹) and counts of hydrogen bonds and salt bridges. Interface contacts were inspected in ChimeraX for residue-level annotation (47).

Per-residue conservation was mapped using ConSurf with homologues selected from the MMseqs2/SeqHub searches and filtered by ConSurf default thresholds for similarity and coverage (48). Final input sets comprised PamA (n = 10) and PamB (n = 9) homologues. ConSurf scores (1→9, 9 = most conserved) were projected onto the experimental models and visualised in ChimeraX; ConSurf default parameters were used.

### Sequence identity matrices

Pairwise sequence identity matrices were generated from full-length alignments produced by Clustal Omega (default settings) (49). Percentage identity values were exported and visualised as heatmaps for figures.

### Particle visualization and micrograph curation

Custom Python scripts were used to preprocess micrographs and overlay particle coordinates for rapid manual inspection (github.com/Luca-Atlas/Micrograph_sorting). Preprocessing steps: sum MRC stacks along z where relevant, twofold upsampling (linear interpolation), Gaussian filtering (σ = 1.0) and percentile-based contrast scaling (0.5–99.5%). Particle coordinates were parsed from RELION STAR files, rescaled to match upsampled images and overlaid as coloured markers to distinguish classes. An interactive Napari (50) workflow allowed sequential review and manual classification of micrographs into different categories via keyboard input; micrographs with <50 particles were pre-filtered.

### Phylogenetic analysis

To survey the distribution of PamA and PamB homologs, we collected GTDB species-representative genomes for candidate phyla radiation (CPR) lineages and a set of non-CPR genomes to show CPR-specific occurrence as used previously (19). The corresponding GTDB phylogenomic tree for CPR was also obtained from GTDB (51).

Putative PamA and PamB homologs were identified using MMseqs2 with an E-value cutoff of 1×10⁻⁵ (52). To improve detection of remote PamA homologs, we additionally used the embedding-based homology search from SeqHub (gLM2 language model) to retrieve candidate sequences with CLIP score > 0.9 (53, 54). The highest scoring language model homolog was chosen as a query in a second MMseqs2 search restricted to CPR genomes.

To evaluate whether language-model–retrieved PamA candidates were structurally consistent with the reference PamA (from M. amalyticus), we performed structure-based comparisons using DALI (55). Specifically, M. amalyticus PamA was aligned to selected candidate PamA structures to assess fold-level similarity (55).

### AlphaFold Predictions

Signal peptides for each protein were predicted using SignalP 6.0 (25) and removed prior to structural prediction. All predictions were conducted using the AlphaFold 3.0 server (alphafoldserver.com) (25). Predicted aligned error (PAE) plots were made using PAE viewer (https://pae-viewer.uni-goettingen.de/) (56).

### Neighborhood analysis

PamA genomic neighborhoods were analyzed for close PamA homologs recovered from the initial MMseqs2 search against GTDB CPR genomes. For each PamA locus, we extracted a ±15 kb window (15 kb upstream and 15 kb downstream) and collected all annotated protein-coding genes within the window. To summarize neighborhood protein diversity and conservation, we constructed a sequence similarity network (SSN) using MMseqs2 similarity scores (edges weighted by bitscore) among neighborhood proteins (n=2188). Markov Cluster Algorithm was used to define clusters and the top 20 most conserved protein-family clusters were identified based on frequency across neighborhoods. Neighborhood-to-neighborhood similarity was then assessed using cblaster and a dendrogram was generated to visualize relationships among PamA neighborhoods (57). The immediate clade that contained the M. amalyticus PamA neighborhood was extracted and visualized as a synteny plot using clinker (58). Genes in the synteny plot were colored according to conserved protein-family membership. To further interpret conserved genes in the PamA neighborhood, we examined proteins annotated as an ATP-binding CBS-domain protein and a multidrug-resistance family (MFS) transporter using predicted structures generated with AlphaFold3 (25). To identify structurally similar proteins, models were queried against the PDB using Foldseek(26), and top hits were selected for structural comparison and visualization in UCSF ChimeraX (26, 44, 47, 59).

## Data visualization

All models and map visualization for figure preparation was conducted within ChimeraX (44, 47), figures were prepared within Inkscape v1.1.

