## Supplementary Information for "A Novel Pilus System in Candidate Phyla Radiation Bacteria"

**Figure S1. Negative-stain electron microscopy of *M. amalyticus* pili:** M. amalyticus cells visualized by negative staining with uranyl acetate. (A) Pilus attached to the *M. amalyticus* cell body. (B–F) Representative images of untethered pili within the same sample.

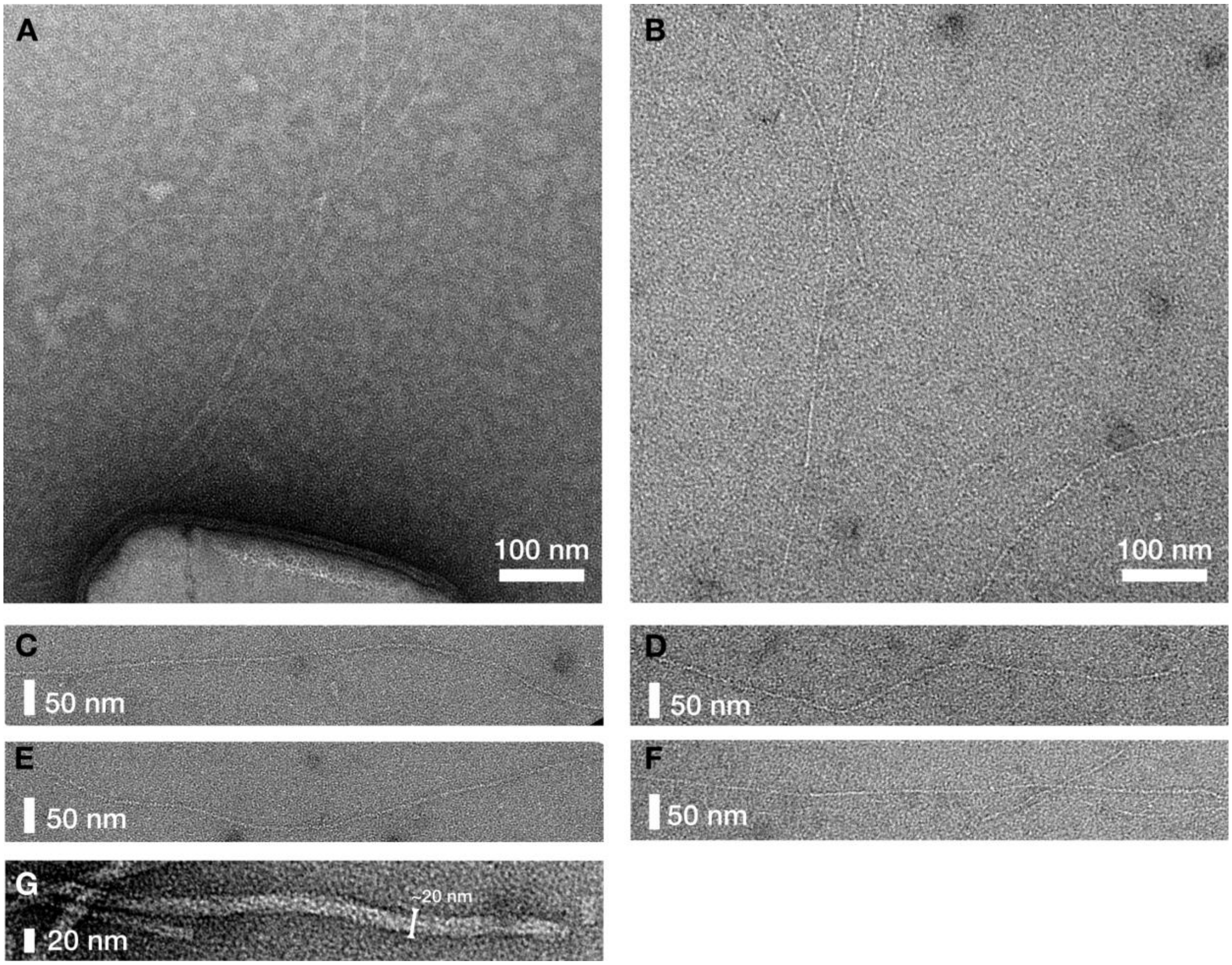

**Figure S2 Single particle cryo-EM data processing.** (A) Initial processing and particle picking of >7,000 movies through MotionCorr2, CTFFIND4 and crYOLO allowed extraction of >1.3 million particles for processing by RELIONv5.0. Left branch: Initially, all particles were roughly aligned through four iterations of RELION 3D classification with a low-resolution reference derived from initially screening data (not shown). The particles were then refined through RELION 2D classification, without any image alignments, and any poorly aligned classes were discarded. The particle alignments were further refined through 3D classification and 3D refinement within RELION and further refined through CTF refinement and Bayesian polishing. Following a final 2D classification of the aligned shiny particles, the final 3D refinement yielded a reconstruction of 2.8 Å resolution composed of 210,417 particles. Right branch: To recover alternate classes of filaments, all picked particles were aligned and classified through 2D classification. A second class of pilins (green box) were selected for further processing. As with the dominant pilus class, particle alignments were refined through 3D classification and 3D refinement, and poorly aligned particles were identified for removal by 2D classification. For the second pilus class, the final 3D refinement yielded a reconstruction of 3.3 Å resolution composed of 150,688 particles.

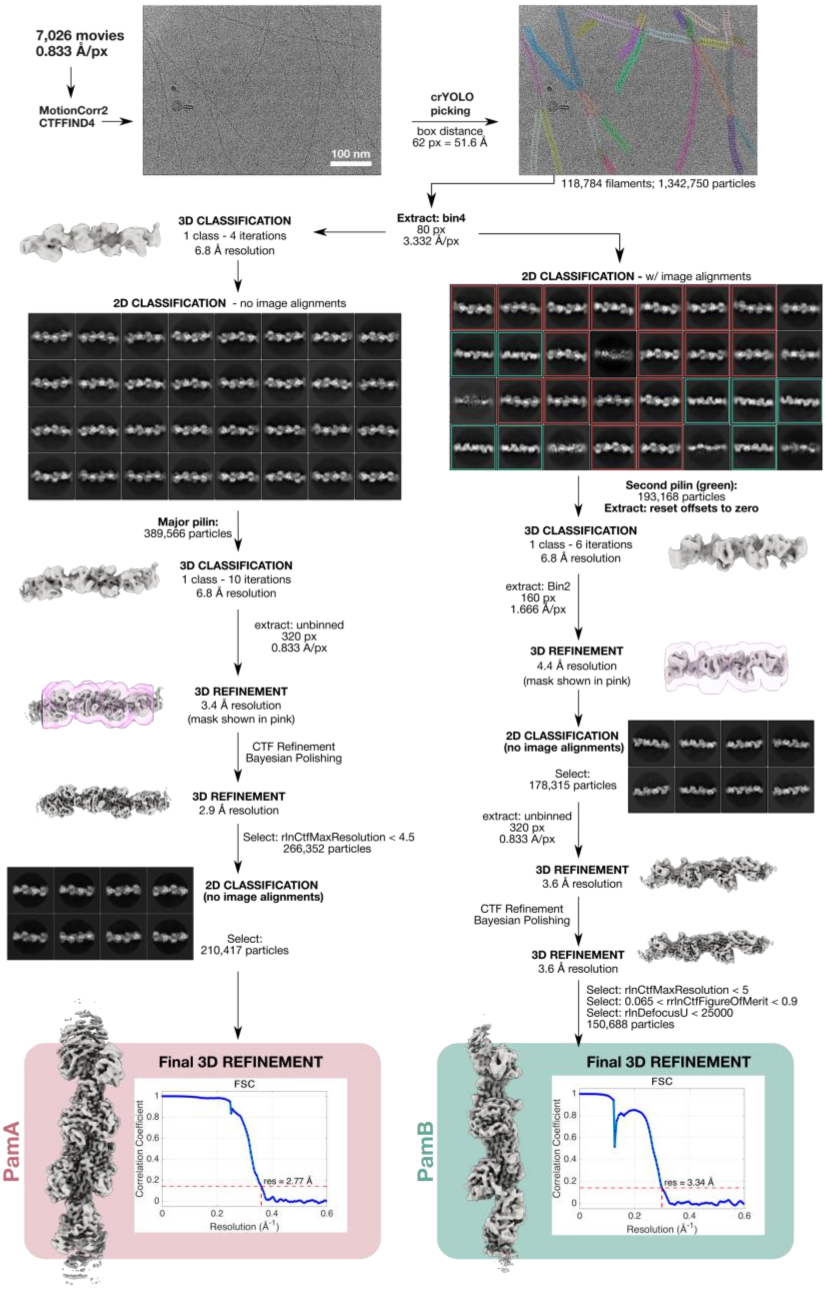

Figure S3: **Model quality for the two Pam pili. (A) The** PamA model PDB: 25PZ / EMD-80296 comprises five PamA chains; three full length pili and two truncated chains. Non-crystallographic symmetry restraints were applied across all five during modelling. Density for the central pilus (chain Aa) is shown within the table and reflects the resolution estimates for the reconstruction (2.8 Å) - local resolution estimates (top right) were calculated within RELION-5.0. (B) The PamB model PDB: 25QA / EMD-80297 comprises five PamB chains; one full length pili and four truncated chains. Non-crystallographic symmetry restraints were applied across all five during modelling. Density for the central pilus (chain A) is shown within the table and reflects the resolution estimates for the reconstruction (3.3 Å) - local resolution estimates (top right) were calculated within RELION-5.0.

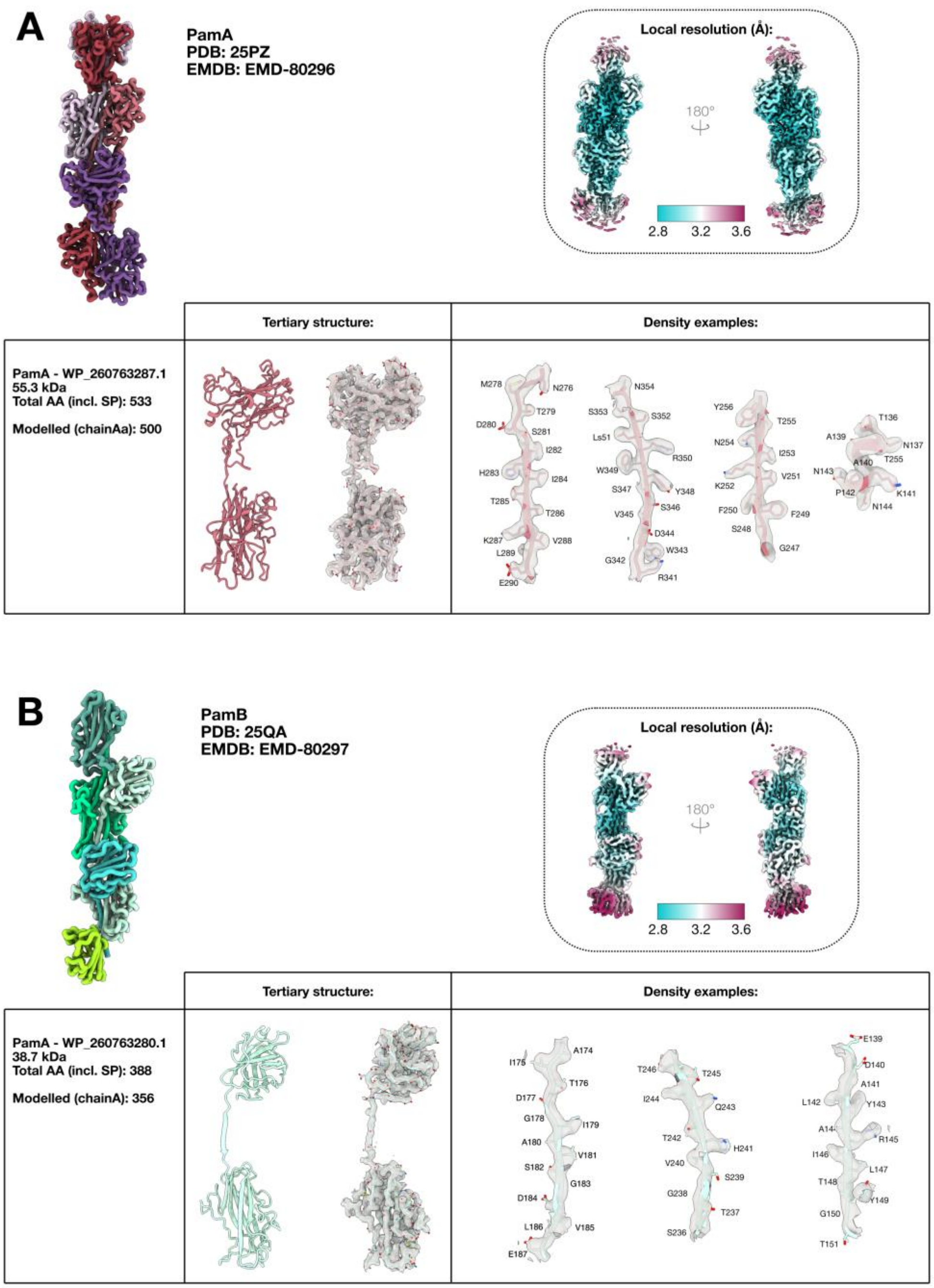

Figure S4 – **PamA and PamB assemble within the same pilus.** Particle coordinates from the final refinements of each pilus class were overlaid onto representative cryo-EM micrographs (PamA, salmon; PamB, green). Multiple clear instances are visible in which contiguous stretches of particles assigned to PamA switch to stretches assigned to PamB within single filaments, consistent with sequential incorporation of different pilin subunits into a single filament.

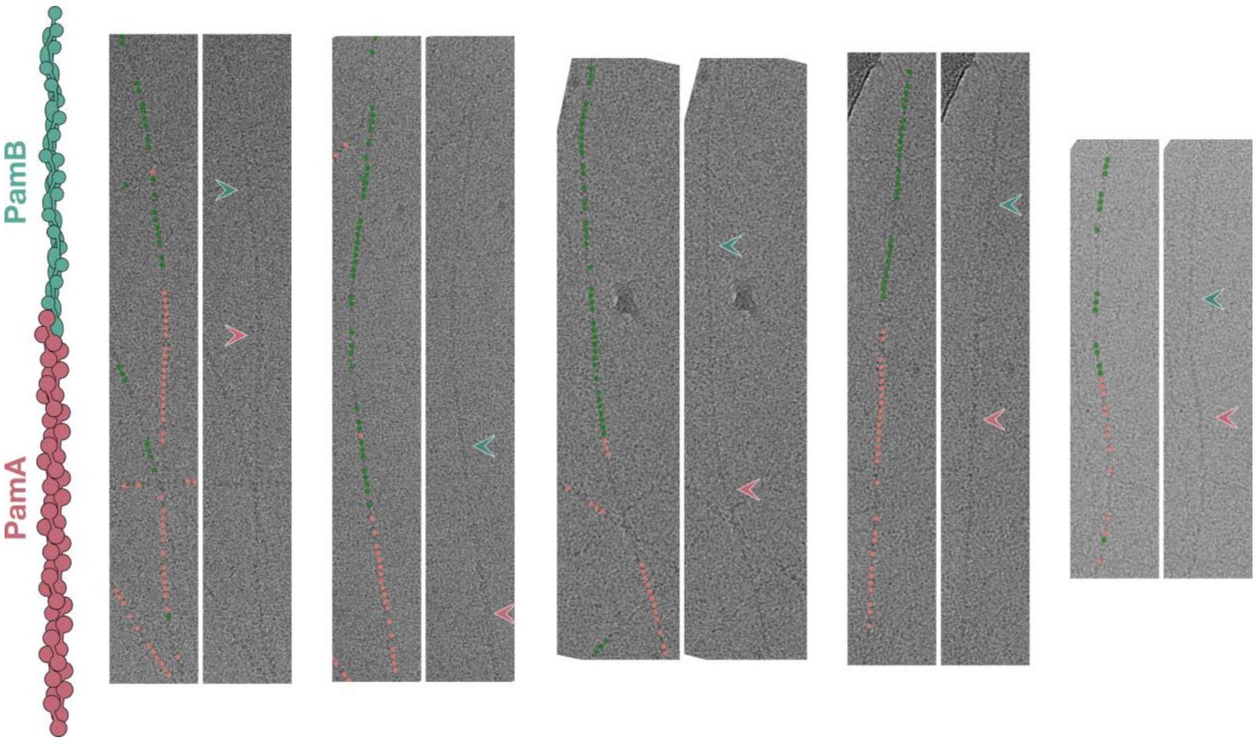

**Figure S5 – Conservation analysis of PamA and PamB (ConSurf).** (A) PamA; (B) PamB. Experimentally determined atomic models are coloured by ConSurf conservation scores calculated from a multiple-sequence alignment of closely related pilins from candidate phyla radiation (CPR) bacteria. Colour scale: maroon = highly conserved, cyan = highly variable (ConSurf scores 9→1). In both pilins, the highest conservation localises to the donated N-terminal linker and the CTD binding site, consistent with a conserved strand-exchange-based assembly mechanism.

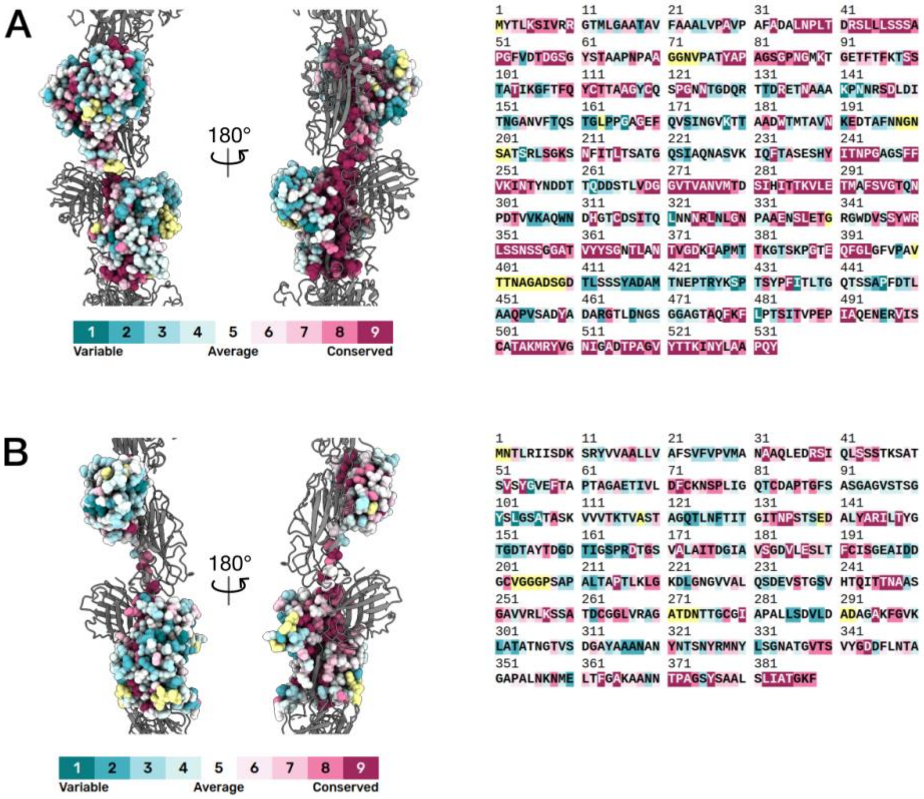

Figure S6 – **Folding, stability and post-translational modifications.** Centre: PamA pilus model shown in surface representation. Right: Magnified view of model with cryo-EM density map at two potential sites for disulphide bond formation within the PamA pilin. Left: Magnified view of Serine 356 within the cryo-EM density map shown with Galactose docked into the extra density observed at this site.

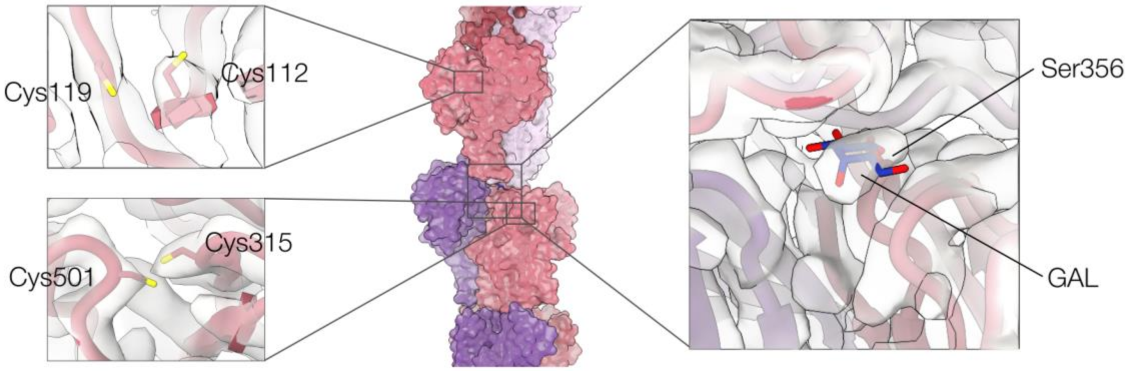

**Figure S7 — Sequence identity and structural homology across Pam pilins.** (A) Pairwise sequence identity matrix for pilins identified in Candidatus Mycosynbacter amalyticus (Saccharimonadia) with a representative PamA structural homolog from Candidatus Roizmanbacteria (Microgenomatia). Values indicate percentage sequence identity derived from full-length alignments; the colour scale reflects identity from low (pale) to high (dark). (B) Structural comparison of all identified M. amalyticus pilins and the Microgenomatia PamA homolog, shown as ribbon models aligned to emphasise the conserved β-barrel architecture of the N-terminal and C-terminal domains. Top: N-terminal domains (NTDs); bottom: C-terminal domains (CTDs). Conserved secondary-structure elements and the overall β-barrel fold are highlighted, illustrating structural conservation despite sequence divergence.

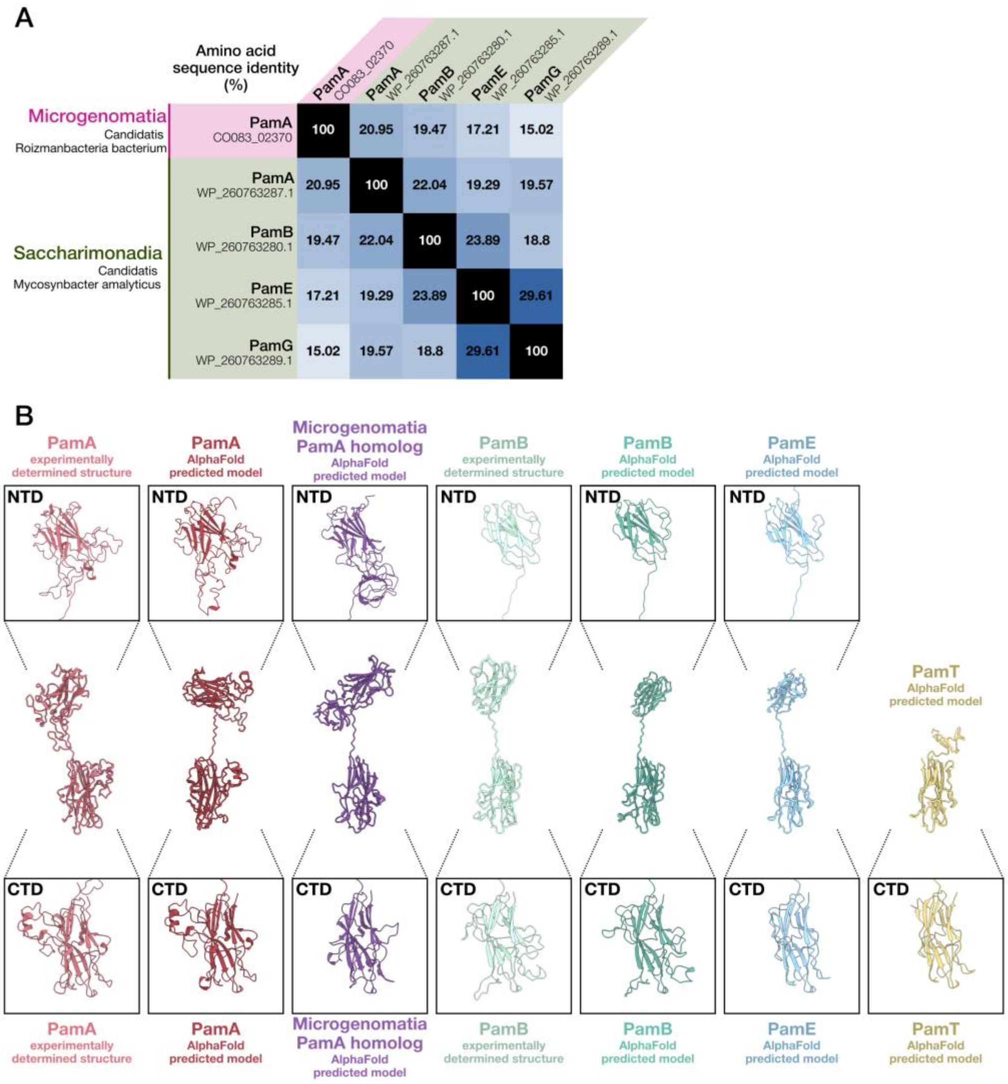

Figure S8 — **AlphaFold3 predictions for pilin–chaperone interactions.**

All five models predicted by AlphaFold3 were overlaid for each pilin–chaperone complex (PamT:PamD; PamB:PamC; PamD:PamA) to illustrate model-to-model variability at the interfaces. Per-residue predicted confidence (pLDDT) is plotted onto the top-ranking model for each prediction and shown as a continuous colour gradient to indicate local model reliability (higher pLDDT = greater confidence). Predicted aligned error (PAE) heatmaps are provided to indicate per-residue positional uncertainty across each model and to highlight regions of inter-chain uncertainty. The predicted template-modelling score (pTM) — reported for each prediction — summarises the overall model confidence and, where applicable, the confidence of inter-chain packing. Together, these panels allow visual assessment of (i) consensus and variability among the five ranked models, (ii) local per-residue confidence, and (iii) positional uncertainty at the pilin–chaperone interfaces.

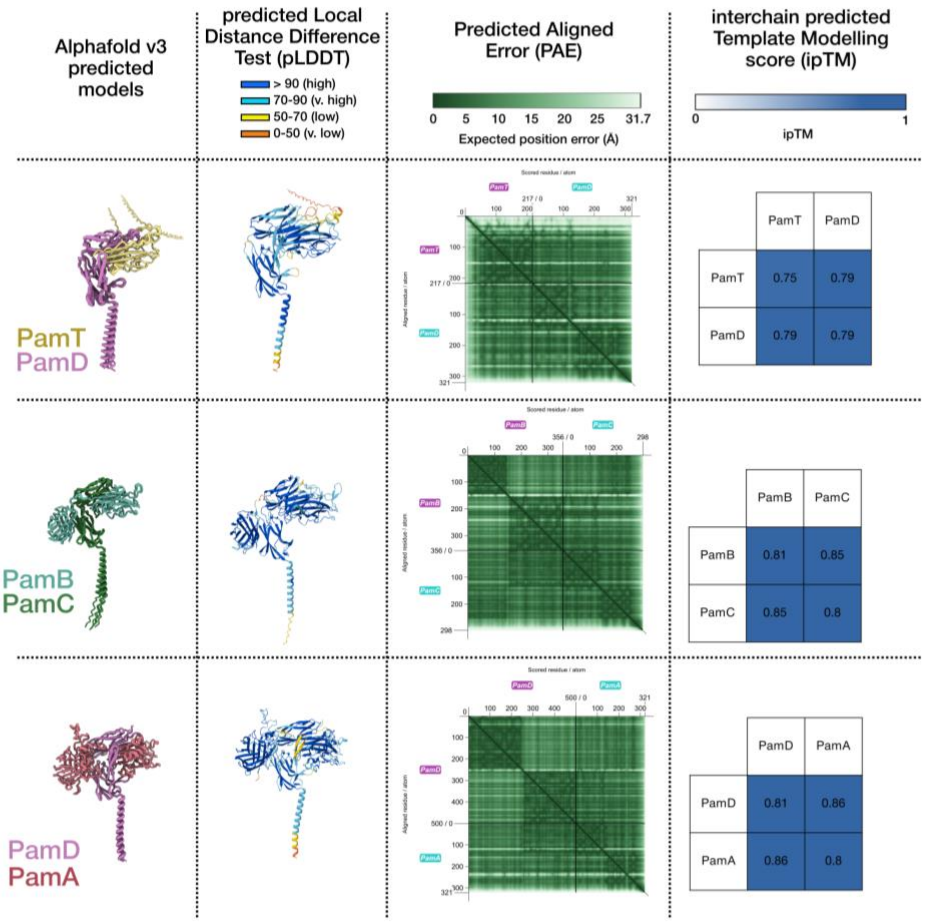

Figure S9 — **AlphaFold3 predictions for Pam pilus assembly complexes.**

Larger combinations of Pam components were combined for prediction of assembly complxes. Top ranked models are shown here with per-residue predicted confidence (pLDDT), Predicted aligned error (PAE) heatmaps and predicted template-modelling score (pTM) — reported for each prediction.

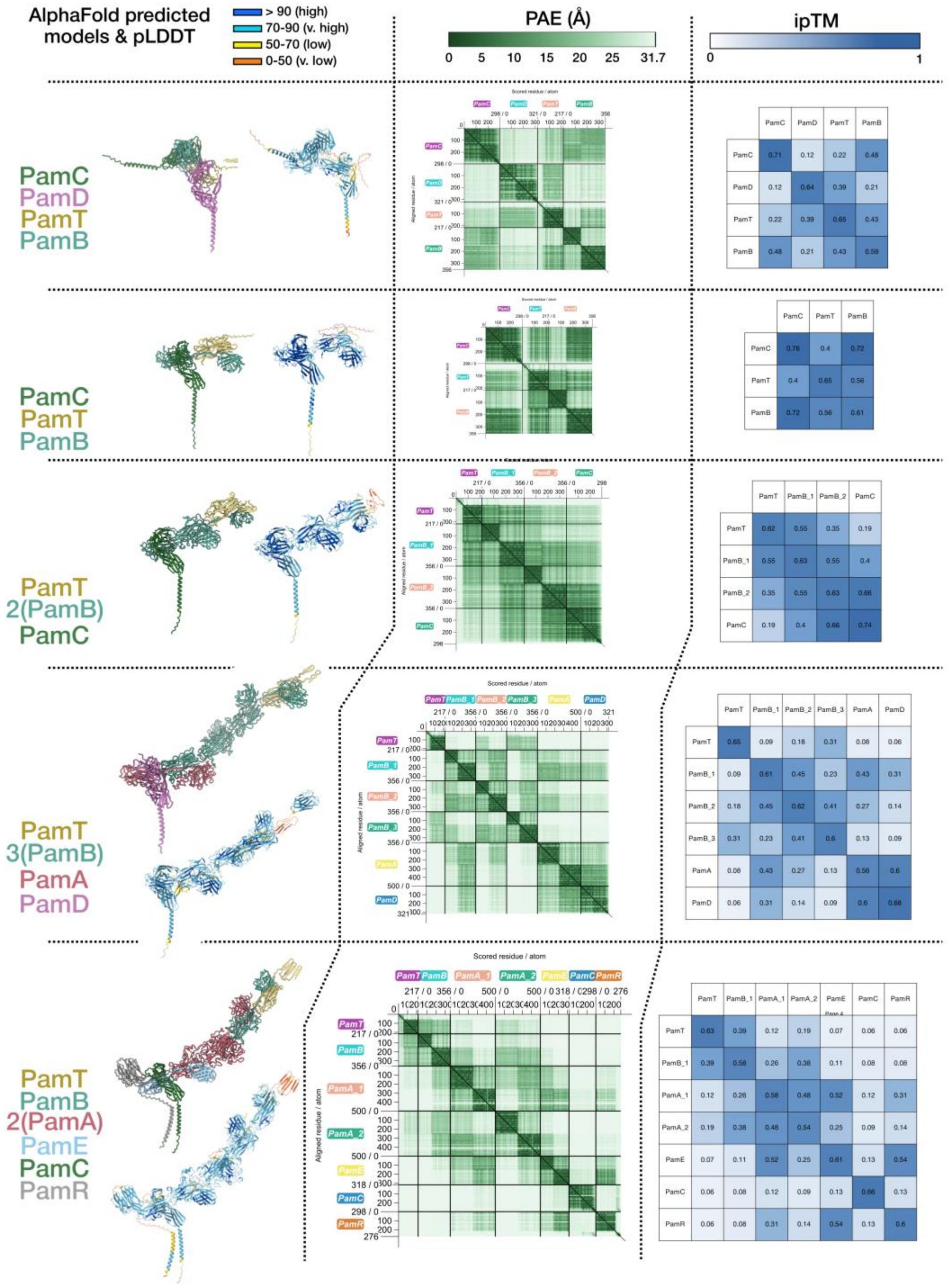

**Figure S10 - AlphaFold3 predictions for Pam neighbourhood transporters showing homology to CorB and MFS families.** (A) Top-ranked AlphaFold3 model for the predicted M. amalyticus CorB homolog (WP_260763263.1). A homodimeric assembly is shown with two putative ATP and two magnesium ions modelled at the nucleotide-binding sites. The panel includes the per-residue pLDDT colouring on the model and the corresponding PAE heatmap (below). For comparison, the inset displays the experimentally determined CorB structure from *Methanoculleus thermophilus* (PDB:7M1T). Structural similarity is evident from the conserved core fold and nucleotide-binding architecture.

(B) Top-ranked AlphaFold3 model for the predicted M. amalyticus major facilitator superfamily (MFS) transporter homolog (WP_260763267.1). The predicted monomeric model is coloured by per-residue pLDDT with the PAE heatmap (below). The inset shows an experimentally determined MFS transporter from *Mycolicibacterium hassiacum* (PDB:8PNL) for structural comparison, highlighting conserved transmembrane helices and the overall 12-helix MFS fold.

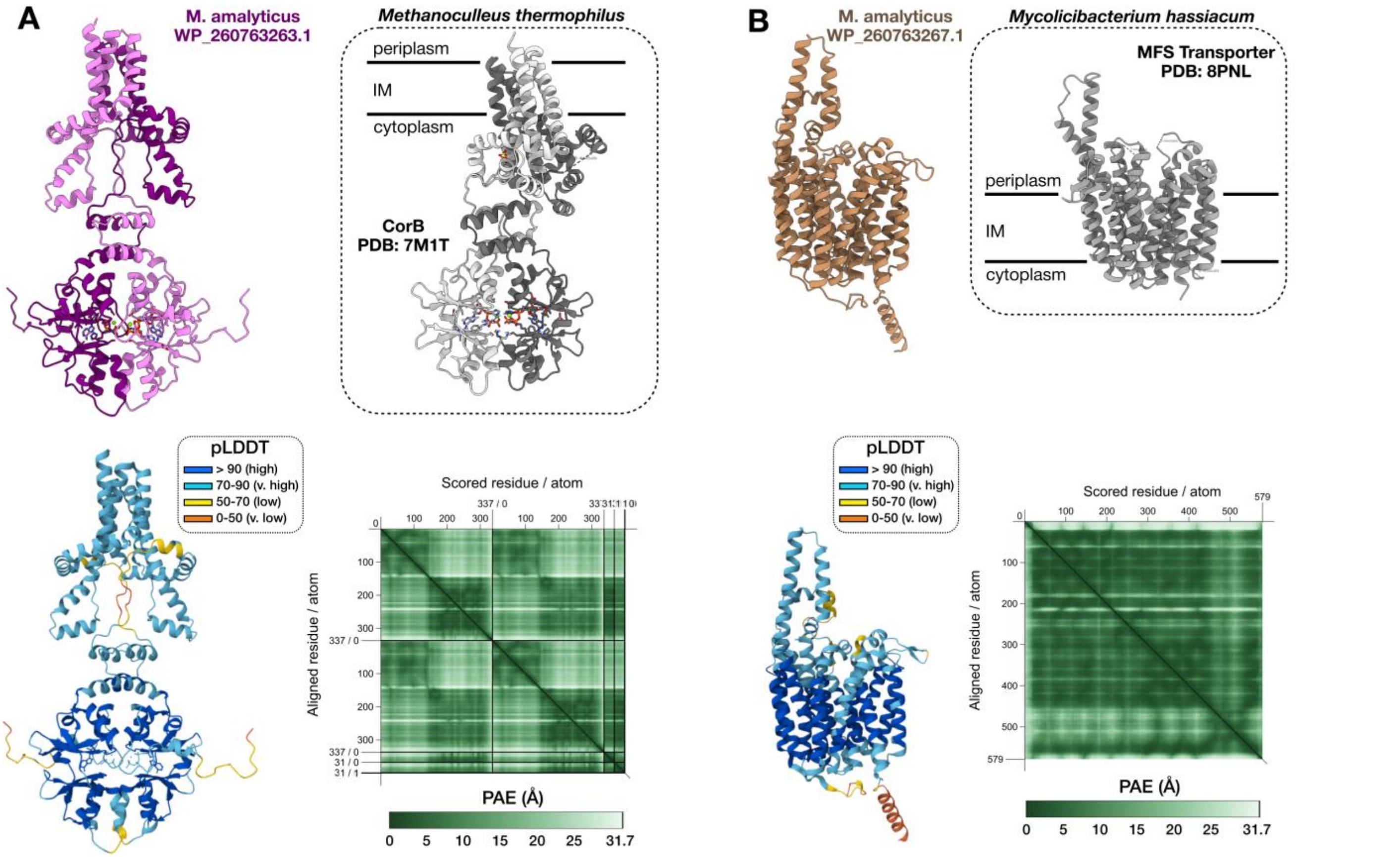

### SUPPLEMENTARY TABLES

Supplementary table 1 – MassSpec results

| Rank | Protein ID | Gene | Protein Description | MW (kDa) | Coverage (%) | # Unique Peptides | Abundances (Normalized) |
|---|---|---|---|---|---|---|---|
| 1 | QHN42848.1 | tuf | Elongation factor Tu | 42.9 | 96 | 47 | 3.54E+11 |
| 2 | QHN43273.1 | pamA | Hypothetical protein (PamA major pilin) | 55.3 | 92 | 31 | 2.07E+11 |
| 3 | QHN42940.1 | gap | Glyceraldehyde-3-phosphate dehydrogenase | 36.2 | 91 | 28 | 1.05E+11 |
| 4 | QHN42861.1 | rpoC | RNA polymerase subunit beta' | 140.1 | 73 | 95 | 8.68E+10 |
| 5 | QHN42862.1 | – | RNA polymerase subunit beta | 123.5 | 71 | 75 | 8.06E+10 |
| 6 | QHN42851.1 | fusA | Elongation factor G | 76.4 | 83 | 56 | 6.34E+10 |
| 7 | QHN43186.1 | dnaK | Molecular chaperone DnaK | 68.5 | 74 | 48 | 4.51E+10 |
| 8 | QHN43237.1 | – | Hypothetical protein | 7.9 | 70 | 9 | 4.45E+10 |
| 9 | QHN43269.1 | pamB | Hypothetical protein (PamB major pilin) | 38.7 | 77 | 20 | 4.30E+10 |
| 10 | QHN43326.1 | – | NAD(P)-binding protein | 59.7 | 76 | 48 | 3.84E+10 |
| 11 | QHN42308.1 | – | Chaperonin GroEL | 56.7 | 86 | 50 | 3.79E+10 |
| 12 | QHN42429.1 | – | Putative pilin | 17.7 | 64 | 12 | 3.78E+10 |
| 13 | QHN42726.1 | atpD | ATP synthase subunit beta | 50.2 | 89 | 34 | 3.63E+10 |
| 14 | QHN42811.1 | – | RNA polymerase subunit alpha | 32.9 | 76 | 26 | 3.60E+10 |
| 15 | QHN42427.1 | pilT | Pilus retraction protein PilT | 39.3 | 89 | 28 | 3.32E+10 |
| 16 | QHN42907.1 | nusA | Transcription termination factor NusA | 41.2 | 87 | 41 | 2.95E+10 |
| 17 | QHN42987.1 | – | LemA family protein | 21.1 | 83 | 20 | 2.91E+10 |
| 18 | QHN42995.1 | – | SDR oxidoreductase | 40.2 | 76 | 25 | 2.44E+10 |
| 19 | QHN42722.1 | – | ATP synthase subunit alpha | 55.3 | 57 | 28 | 2.42E+10 |
| 20 | QHN42474.1 | – | S1 RNA-binding protein | 38.6 | 74 | 27 | 2.40E+10 |
| 21 | QHN42869.1 | tig | Trigger factor | 47.1 | 78 | 44 | 2.39E+10 |
| 22 | QHN42271.1 | – | Hsp20 family protein | 17.4 | 22 | 5 | 2.26E+10 |
| 23 | QHN42909.1 | – | Fructose-bisphosphate aldolase class I | 72.1 | 70 | 34 | 2.18E+10 |
| 24 | QHN43208.1 | hflB | ATP-dependent zinc metalloprotease FtsH | 68.2 | 69 | 50 | 2.00E+10 |
| 25 | QHN43182.1 | – | FAD-binding protein | 62.8 | 68 | 33 | 1.96E+10 |
| 26 | QHN43051.1 | – | 50S ribosomal protein L25 | 25.4 | 84 | 15 | 1.96E+10 |
| 27 | QHN42422.1 | pilM | Type IV pilus assembly protein PilM | 38 | 61 | 20 | 1.85E+10 |
| 28 | QHN42542.1 | secA | Preprotein translocase subunit SecA | 99.5 | 63 | 59 | 1.78E+10 |
| 29 | QHN42381.1 | – | Hypothetical protein | 71.6 | 45 | 23 | 1.78E+10 |
| 30 | QHN43271.1 | pamD | Hypothetical protein (PamD pilus chaperone) | 37.4 | 67 | 22 | 1.77E+10 |
| 31 | QHN42555.1 | – | Hypothetical protein | 40.5 | 71 | 30 | 1.77E+10 |
| 32 | QHN43322.1 | rfbA | Glucose-1-phosphate thymidylyltransferase | 30.7 | 71 | 15 | 1.77E+10 |
| 33 | QHN42522.1 | rpsB | 30S ribosomal protein S2 | 25.5 | 79 | 18 | 1.76E+10 |
| 34 | QHN42624.1 | – | Response regulator | 27.9 | 45 | 12 | 1.76E+10 |
| 35 | QHN43198.1 | – | Alpha/beta hydrolase domain protein | 61.5 | 65 | 26 | 1.69E+10 |
| 36 | QHN42821.1 | rpsE | 30S ribosomal protein S5 | 21.2 | 59 | 12 | 1.69E+10 |
| 37 | QHN42426.1 | pilB | Type IV pilus assembly protein PilB | 64 | 77 | 35 | 1.69E+10 |
| 38 | QHN43268.1 | – | Hypothetical protein | 36.5 | 66 | 21 | 1.65E+10 |
| 39 | QHN42421.1 | pamC | Hypothetical protein (PamC pilus chaperone) | 119 | 58 | 49 | 1.64E+10 |
| 40 | QHN42844.1 | rpsJ | 30S ribosomal protein S10 | 11.4 | 84 | 15 | 1.63E+10 |
| ... | ... | ... | ... | ... | ... | ... | ... |
| 631 | QHN43270.1 | pamR | Hypothetical protein (PamR pilus tether) | 32.8 | 5 | 1 | 2.20E+08 |
| 651 | QHN43272.1 | pamE | Hypothetical protein (PamE minor pilin) | 38.6 | 30 | 5 | 1.81E+08 |
| 768 | QHN43274.1 | pamT | Hypothetical protein (PamT tip pilin) | 25.6 | 44 | 4 | 4.23E+07 |

**Supplementary Table 2 – CryoEM**

| | **Single particle analysis** | |
|---|---|---|
| magnification | 105, 000 x | |
| voltage (keV) | 300 | |
| electron exposure ($e^{-}/Å^2$) | 55.8 | |
| defocus range | -0.5 to -2.4 | |
| pixel size (Å) | 0.833 | |
| symmetry imposed | C1 | |
| movies | 7026 | |
| particle picking method | crYOLO | |
| box distance (px) | 62 | |
| minimum filament length (particles) | 4 | |
| threshold | 0.3 | |
| box size (px) | 448 | |
| initial particles (total) | 1,342,750 | |
| **ARCHITECTURE** | **Class 1 - PamA** | **Class 2 - PamB** |
| initial particles (after 2D classification) | 389,566 | 193,168 |
| final particles | 210,417 | 150,688 |
| **Full map resolution (Å)** | 2.77 | 3.34 |
| FSC threshold | 0.143 | 0.143 |
| **REFINEMENT** | | |
| sharpening factor | n/a | n/a |
| map sharpening method | deepEMhancer | deepEMhancer |
| model resolution | 2.6 | 2.7 |
| FSC threshold | 0.143 | 0.143 |
| **MODEL COMPOSITION** | | |
| chains | 5 | 5 |
| non-hydrogen atoms | 28461 | 14620 |
| protein residues | 1999 | 1068 |
| **B FACTORS ($Å^2$)** | | |
| protein | 165.02 | 96.54 |
| nucleotide | - | - |
| ligand | - | - |
| **BONDS (RMSD)** | | |
| bond length (Å) | 0.002 | 0.002 |
| bond angles (°) | 0.430 | 0.403 |
| **VALIDATION** | | |
| MolProbity score | 0.81 | 0.83 |
| Clashscore, all atoms: | 1.09 | 1.16 |
| poor rotamers (%) | 1.14 | 3.83 |
| favored rotamers (%) | 98.86 | 96.17 |
| **RAMACHANDRAN PLOT** | | |
| Ramachandran outliers (%) | 0.00 | 0.00 |
| Ramachandran favoured (%) | 98.74 | 98.77 |

| | | |
|---|---|---|
| Rama distribution Z-score | 0 | -0.83 |
| **DEPOSITION** | | |
| EMDB | EMD-80296 | EMD-80297 |
| PDB | 25PZ | 25QA |
